# scMTG reconstructs single-cell temporal dynamics with Markov transition generators

**DOI:** 10.64898/2026.06.04.730241

**Authors:** Xuejian Cui, Haochen Wang, Zijing Gao, Rui Jiang, Wing Hung Wong, Qiao Liu

**Affiliations:** Ministry of Education Key Laboratory of Bioinformatics, Bioinformatics Division at the Beijing National Research Center for Information Science and Technology, Center for Synthetic and Systems Biology, Department of Automation, Tsinghua University, Beijing 100084, China; Department of Biostatistics, Yale University, New Haven, CT 06511, USA; Department of Statistics, Stanford University, Stanford, CA, 94305, USA; Department of Biomedical Data Science, Stanford University, Stanford, CA, 94305, USA; Program in Computational Biology and Bioinformatics, Yale University, New Haven, CT 06511, USA

## Abstract

Single-cell time-series data offer a unique opportunity to study how cell states emerge, evolve, and diversify over time. However, cells are destructively measured across time points and individual cells cannot be directly tracked, making it challenging to reconstruct cell-state transitions and uncover the dynamic regulatory programs. Existing methods are predominantly based on optimal transport and typically require predefined low-dimensional representations, which can limit scalability, flexibility, and mechanistic interpretability. Here we present scMTG, a single-cell Markov transition generative framework that jointly learns cell representations and temporal cell-state transitions from unpaired single-cell time-series data. In a series of experiments, scMTG demonstrates superior performance in interpolating held-out missing-time-point data and inferring cross-time-point transition matrices, compared with state-of-the-art methods. More importantly, the learned Markov transition generator provides an interpretable framework for identifying the molecular programs that drive cell-state changes, enabling characterization of cell fate decisions and construction of time-resolved regulatory networks. Together, scMTG provides a unified generative framework for reconstructing cellular dynamics and uncovering regulatory programs that shape development, differentiation, and disease progression. scMTG is available as open-source software at https://github.com/liuq-lab/scMTG.

## Introduction

Single-cell sequencing technologies have revolutionized the study of gene regulatory mechanisms at the resolution of individual cells^1-6^. However, conventional single-cell sequencing primarily captures static snapshots of specific differentiation states, whereas most biological processes, including cell differentiation, tissue development, and disease progression, are inherently dynamic^7, 8^. By enabling single-cell resolution with temporal information, single-cell time-series data allow researchers to examine the dynamic changes in molecular signals across multiple time points, laying the foundation for investigating the molecular mechanisms underlying dynamic biological processes^9-12^.

Since single-cell sequencing technologies are inherently destructive for cells, single-cell time-series data are typically obtained from different batches of cells at different time points, with no direct correspondence between cells across time points^7, 13-17^. Therefore, establishing transition relationships between cells at different time points, identifying key regulatory genes and constructing temporal gene regulatory networks (GRNs) that drive dynamic biological processes, remain both highly challenging and critically important in the analysis of single-cell time-series data^7, 18^.

Recent advances in computational methods have shown strong potential for time-series single-cell analysis^7, 13, 18-21^. For example, Waddington-OT (WOT) formulates the transition relationships between different cell populations at adjacent time points as an optimal-transport (OT) problem, thereby recovering ancestor-descendant fates based on the transport maps^7^. CellRank 2 integrates intra-time-point cell state transitions computed by CellRank^22^ with inter-time-point transitions estimated using WOT^7^ or moscot^18^ to construct a final transition matrix, enabling a wide range of downstream temporal analyses. moscot is a scalable OT-based framework for mapping cells across temporal and spatial samples, and has demonstrated strong performance in applications such as reconstructing mouse embryogenesis and resolving lineage trajectories during mouse pancreas development^18^. PRESCIENT represents a complementary generative approach that formulates differentiation as a stochastic diffusion process in a low-dimensional cell-state space. It learns a global neural potential landscape whose negative gradient defines the drift of cell-state trajectories, enabling held-out time-point reconstruction and fate prediction through forward trajectory simulation^19^.

However, existing methods generally suffer from several limitations when modeling and characterizing single-cell time-series data. First, current leading approaches are largely based on OT^7, 13, 18^, which estimates discrete transport plans or transition matrices between empirical cell populations at adjacent time points. While such matrices are useful for matching cell distributions and inferring ancestor-descendant relationships, they remain static, sample-level summaries of transitions between observed cells. They do not explicitly learn the general transition function that describes how a molecular state evolves over time. Second, current computational frameworks rely on multi-stage pipelines in which representation learning, transition inference, and downstream fate analysis are performed separately. For example, they often used predefined low-dimensional representations derived from dimensionality reduction techniques such as principal component analysis (PCA) or latent semantic indexing (LSI)^7, 13, 18, 19^. Such separation may make the inferred transitions sensitive to preprocessing choices and may limit the flexibility of the model in capturing complex temporal dynamics. Third, since the primary output of these methods is transition matrices, low-dimensional trajectories, or fate probabilities, rather than differentiable gene-level transition functions, temporal regulatory analyses are usually performed post hoc. For example, regulatory interpretation is commonly based on inferred cell-state transitions, fate probabilities, or simulated trajectories. These limitations highlight the critical need for a framework that explicitly learns generative and differentiable transition functions from unpaired single-cell time-series data, thereby enabling cell state transition inference and dynamic regulatory mechanism discovery within a unified model-based framework.

To address these gaps, we develop scMTG, a single-cell Markov transition generative modeling framework for learning cell state transitions and dynamic gene regulatory programs from unpaired single-cell time-series data. scMTG integrates a parameter-shared autoencoder with time-indexed Markov transition generators, thereby jointly learning a shared low-dimensional representation across time points and a series of conditional generative transition functions between adjacent time points. To account for changes in cell population composition caused by proliferation and apoptosis, scMTG further incorporates a cell-specific growth-weighted sampling strategy during transition learning. Unlike OT-based methods that primarily infer static transport matrices, scMTG explicitly learns transition functions that map the cell state to its future state distribution. These learned transition kernels can be used to infer cross-time-point transition matrices, interpolate held-out intermediate time points, and characterize lineage relationships and fate-related temporal dynamics. More importantly, since the Markov transition generators are continuous and differentiable, scMTG enables direct characterization of how gene expression at one time point influences gene expression at the next time point, providing a model-based route to identify key regulatory transition-driving genes and construct temporal and cell-type-specific GRNs directly from the learned transition functions. Across multiple benchmark datasets, scMTG outperforms existing methods in recovering held-out intermediate time points and inferring biologically meaningful transition relationships, while also revealing dynamic regulatory programs underlying cell fate decisions. In summary, scMTG shifts single-cell temporal analysis from estimating static couplings of adjacent time points toward learning differentiable temporal transition functions, providing a unified framework for modeling cellular dynamics, lineage progression, and regulatory mechanisms in cell development, differentiation, and disease progression.

## Results

### Overview of the scMTG framework

scMTG is a Markov transition generative framework to model cell state transition and dynamic regulation analysis for single-cell time-series data. Taking the processed cell-by-gene matrices of single-cell time-series data as input, scMTG first employs a parameter-shared autoencoder module to generate low-dimensional representations of cells at different time points, and subsequently utilizes multiple Markov transition generators to model cell state transitions between adjacent time points (**Fig. 1**a and Methods). We adopt adversarial training to learn the Markov transition generators across different time points. Specifically, within the Markov transition module of scMTG, the generator synthesizes cells at the current time point based on cells from the previous time point, while the discriminator distinguishes between generated and real cell profiles. After model training, the generator acquires the ability to model the cell state transitions between consecutive time points. To account for cell proliferation and apoptosis^7, 18, 19^, we adopt an estimated cell-specific growth-weighted sampling strategy during the training process (see Methods).

**Fig. 1.**
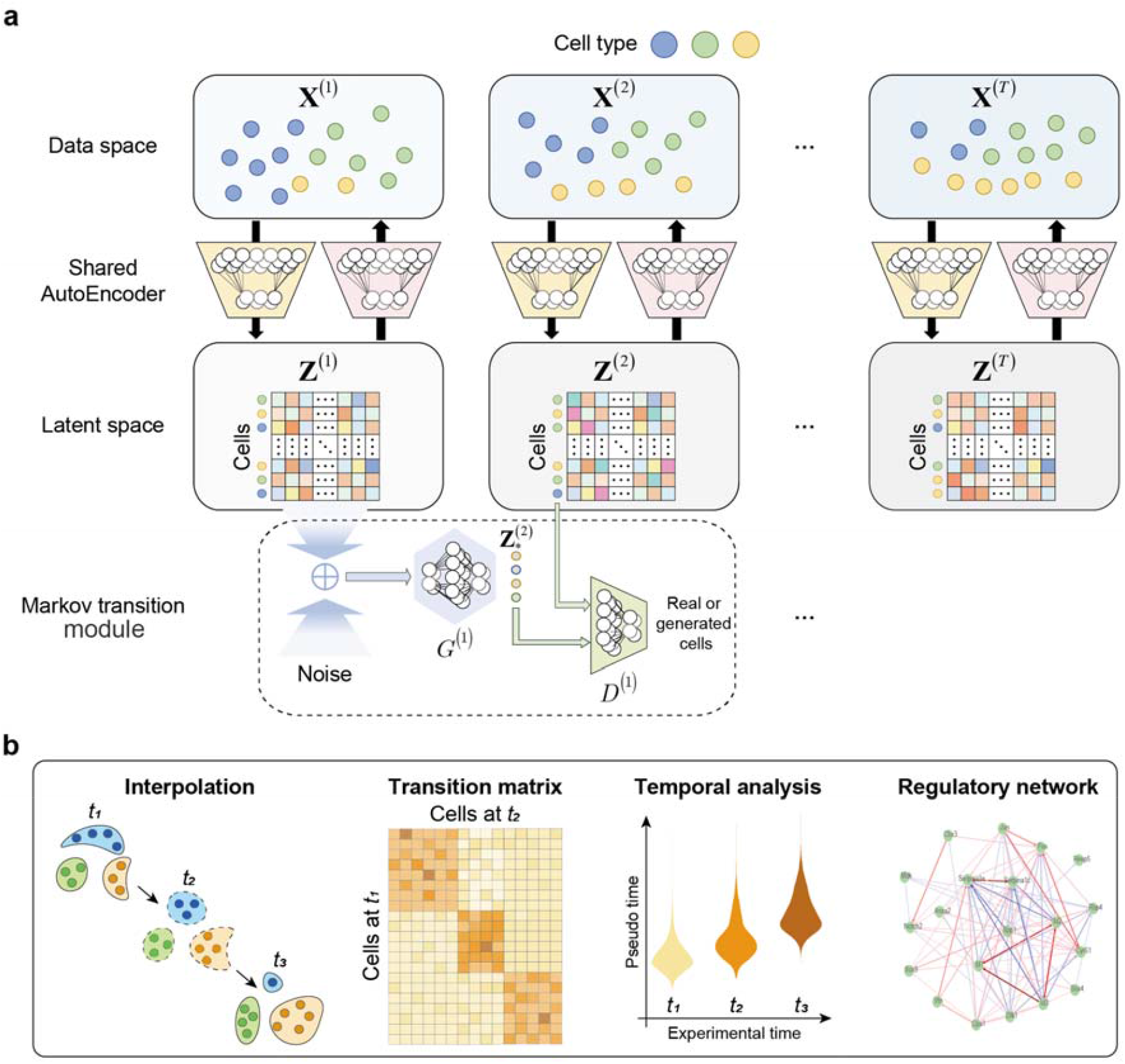
Overview of scMTG. **a**, The architecture of scMTG model. scMTG is a deep generative model that consists of an autoencoder module and a Markov transition module. The autoencoder module of scMTG is parameter-shared across all time points. The Markov transition module of scMTG models cell state transitions between adjacent time points using generative adversarial networks. **b**, scMTG facilitates data interpolation, generation of transition matrix, temporal analysis and construction of regulatory network.

Leveraging the learned cell embeddings and cell state transitions, scMTG 1) enables accurate interpolation of missing time points, 2) performs effective inference of transition matrices across time points, 3) facilitates interpretable temporal analysis, and 4) enables the construction of temporal regulatory networks and precise elucidation of dynamic regulatory mechanisms (Fig. 1b). Overall, these capabilities make scMTG a powerful tool for single-cell temporal analysis and provide important insights into the molecular mechanisms underlying dynamic biological processes.

### scMTG accurately reconstructs held-out temporal cell-state distributions

To systematically assess the capability of scMTG to interpolate single-cell data of missing time points, we conducted experiments aimed at recovering held-out cells of intermediate time points. Specifically, scMTG was benchmarked on 6 single-cell time-series datasets (Supplementary Table 1) against four baseline methods, including WOT^7^, PRESCIENT^19^, CellRank 2^13^ and moscot^18^, using evaluation metrics including maximum mean discrepancy (MMD)^23, 24^, Spearman’s correlation coefficient (SCC)^18^, and local inverse Simpson’s index (LISI)^25^ (see Methods).

We benchmarked scMTG against four baseline methods across interpolation tasks spanning multiple intermediate time points on 6 benchmark datasets. scMTG outperforms the baseline methods by achieving the overall best interpolation performance and exhibiting strong stability across multiple time points (**Fig. 2**a and Supplementary Fig. 1). Taking the Serum dataset as an example^7, 13^, scMTG achieves a significant improvement of 12.2% in median LISI (one-sided paired Wilcoxon signed-rank tests p-values < 8e-7) and 4.3% in median SCC (p-values < 8e-8), compared with the second-best method. Across the interpolation tasks of 37 intermediate time points, scMTG obtains the best LISI values in 34 time points (91.9%), the best MMD values in 32 time points (86.5%), and the best SCC values in 35 time points (94.6%) (Fig. 2b and Supplementary Fig. 2).

**Fig. 2.**
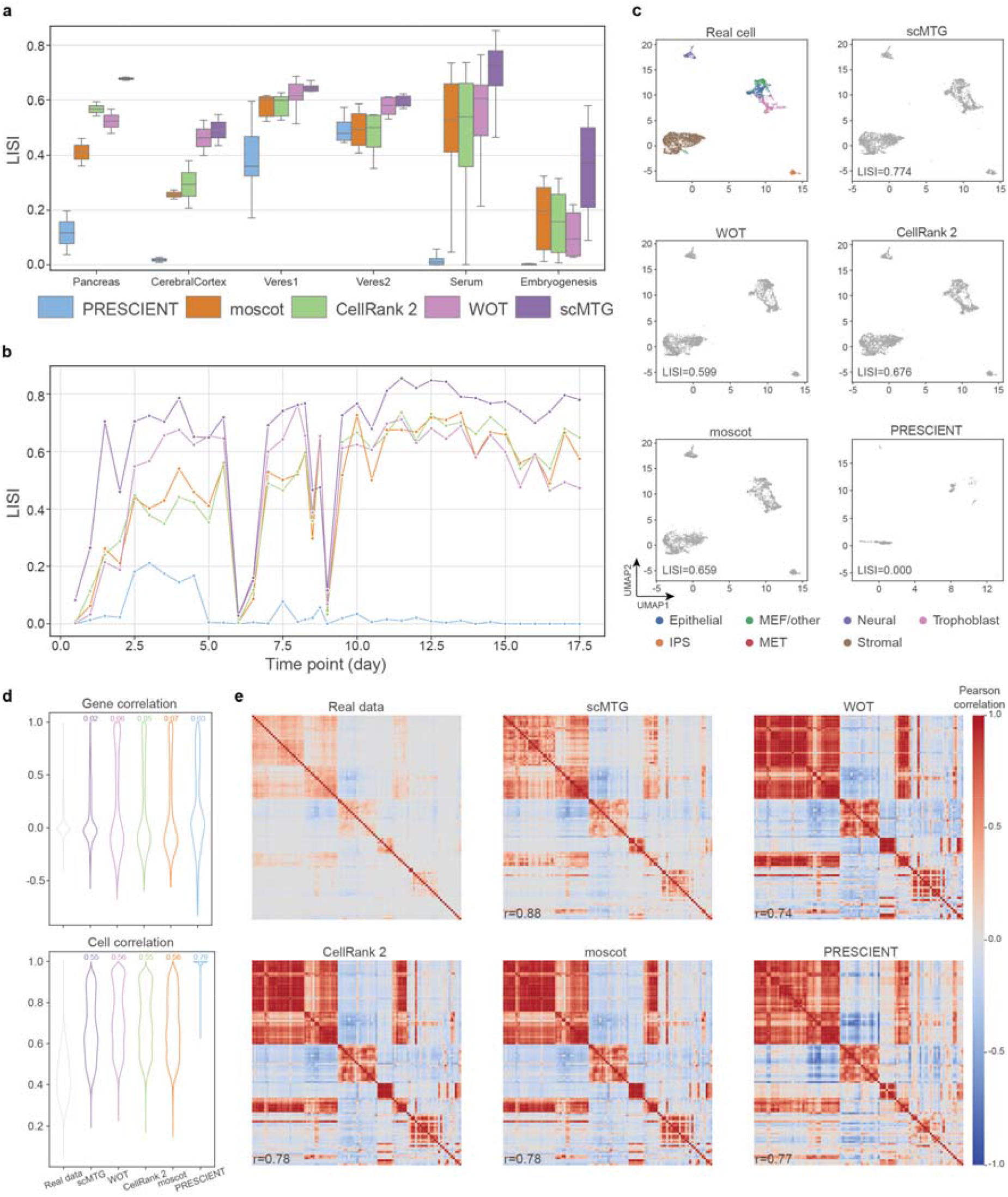
scMTG interpolates single-cell data of missing time points. **a**, Interpolation results of different methods evaluated by LISI on 6 benchmark datasets. **b**, Interpolation results of different time points evaluated by LISI on the Serum dataset. **c**, UMAP visualization of the real cells on Serum dataset, compared with the generated cells derived from different methods. **d**, Distributions of gene correlation (top) and cell correlation (bottom) in the real data and generated data from different methods on the Serum dataset. The number on top of each violin plot is the Kolmogorov-Smirnov (KS) distance between the generated data distribution and the real data distribution. **e**, Heatmaps of the gene-gene correlation matrices (showing top 100 highly expressed genes) in the real data and the generated data from different methods on the Serum dataset.

To further evaluate the distributional consistency between the generated and real cells, we conducted a series of comparative experiments on the Serum dataset^7, 13^. First, as shown in the UMAP visualization at the “day 15.0” time point, the generated cells by scMTG exhibit highly consistent distribution to the held-out real cells, reflected by the highest LISI value (Fig. 2c). Specifically, the generated cells by scMTG are seamlessly integrated with real cells and faithfully preserve the cell heterogeneity at the intermediate time point (Supplementary Fig. 3e). In contrast, the generated cells by PRESCIENT (highlight with blue circle) and moscot (highlight with orange circle) exhibit evident batch effects relative to real cells (Supplementary Fig. 3a-b). CellRank 2 struggles to illustrate a clear separation between cells in green circle (Supplementary Fig. 3c). The generated cells by WOT contain an aberrant outlying cell population (highlight with pink circle) (Supplementary Fig. 3d). Second, referring to scDesign3^26^, we displayed the distributions of gene-gene correlations for the real data and the generated data by different methods, and computed the Kolmogorov-Smirnov (KS) distance between the distributions of real and generated data (Fig. 2d). Compared with baseline methods, the distributions of gene-gene correlation between the generated data by scMTG and real data showed the strongest concordance, reflected by the smallest KS distance. Similarly, scMTG outperformed baseline methods in generating more realistic cells and in better preserving the cell-cell correlations, reflected by smaller KS distance (Fig. 2d). Third, we constructed the gene-gene correlation matrices for the top 100 highly expressed genes in the real data and the generated data by different methods, and quantified the similarity of generated data to the correlation matrix of real data using the Pearson’s correlation coefficient (Fig. 2e). The correlation matrix of generated data by scMTG achieves the highest similarity to that of the real data than baseline methods.

Collectively, these results demonstrate that scMTG excels in both accurate temporal interpolation and informative visualization, highlighting its capacity to reveal temporal dynamics and cell heterogeneity.

### scMTG effectively infers transition matrix across time points

Given the unpaired nature of cells across different time points in single-cell time-series data, it is crucial for temporal analysis to infer the transition matrix that captures differentiation relationships between time points^7, 13, 18^. Based on the generator function within the Markov transition module of scMTG, we obtained a transition matrix that characterizes the transition probabilities between cells at adjacent time points (see Methods). To further validate the biological significance of the transition matrix, we aggregated it to the cell-type level, thereby constructing a matrix of transition probabilities among cell types between adjacent time points.

We evaluated the performance and interpretability of the transition matrix generated by scMTG on the Embryogenesis dataset^16^, which benefits from the well-established differentiation hierarchies and lineage relationships among cell types^16, 18, 27, 28^. The transition matrices generated by scMTG uncover the differentiation lineage, which is clearly consistent with the mouse embryonic development process (Fig. 3a and Supplementary Fig. 4). Moreover, we introduced two evaluation metrics defined at the germ-layer and cell-type levels (see Methods), and observed that scMTG obtains significant higher germ-layer transition accuracy (one-sided paired Wilcoxon signed-rank tests p-values < 4e-2) than baseline methods except WOT (p-value = 0.07), and significant higher cell-type transition accuracy (one-sided paired Wilcoxon signed-rank tests p-values < 2e-2) than baseline methods. For example, scMTG achieves accurate germ-layer transitions (highlighted by green squares in Fig. 3a), such as the transition from “Caudal mesoderm” to “Extraembryonic mesoderm”, and cell-type transitions (highlighted by blue squares in Fig. 3a), such as the transition from “Hematoendothelial progenitors” to “Endothelium”, both of which are supported by existing literature^16, 18, 27, 28^ (Supplementary Table 2).

**Fig. 3.**
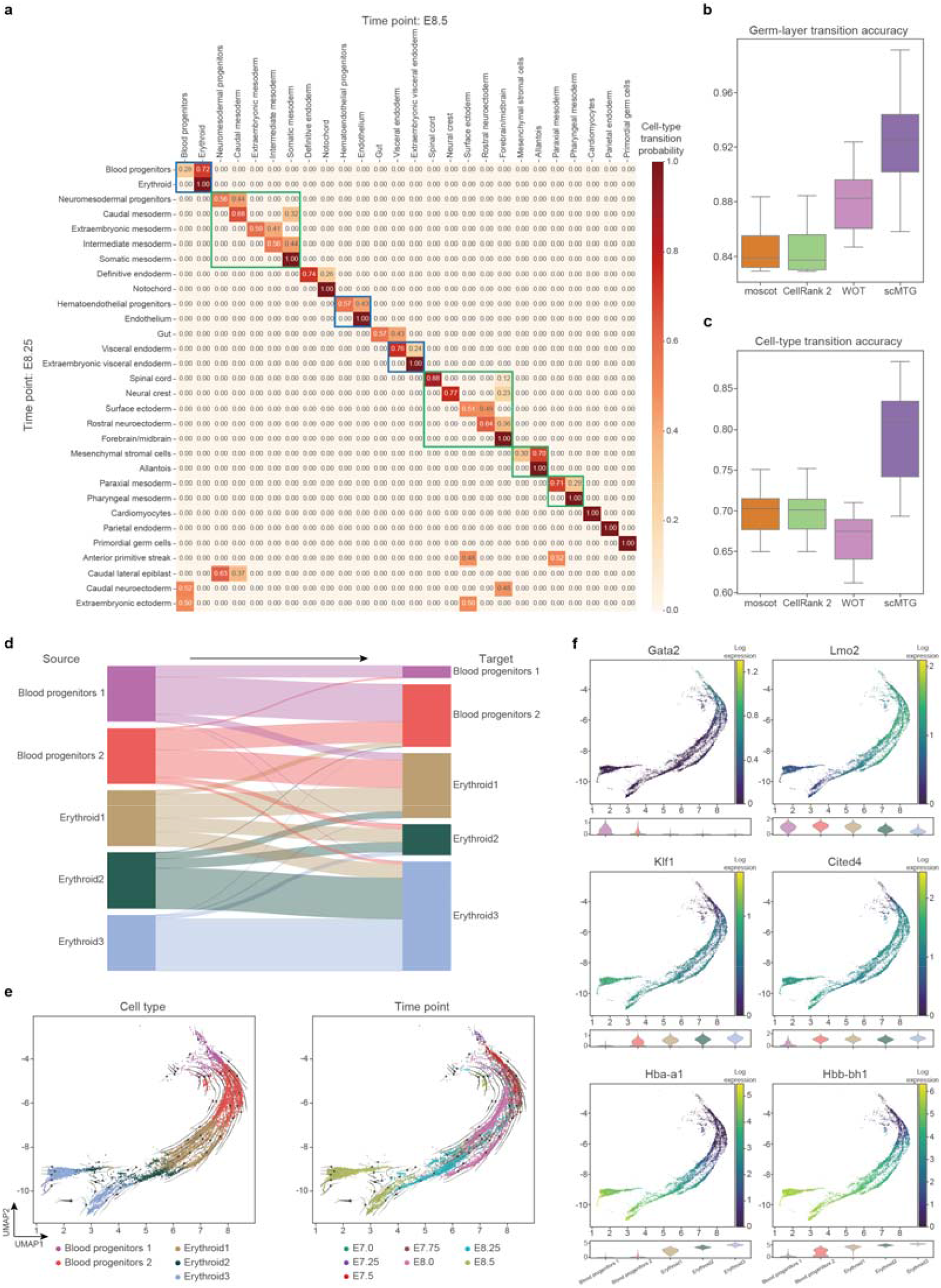
scMTG provides transition matrix of adjacent time points. **a**, Heatmap visualizing transition probabilities of cell types from E8.25 to E8.5 on the Embryogenesis dataset as obtained using scMTG. **b-c**, Accuracy comparison between scMTG and baseline methods in terms of germ-layer (**b**) and cell-type transition scores (**c**) by developmental stage. **d**, Descendancy probabilities of cell types in erythropoiesis on the Embryogenesis dataset as obtained using scMTG. **e**, UMAP visualization of erythropoiesis on the Embryogenesis dataset with inferred velocities by CytoTRACE, colored by cell types (left) and time points (right). **f**, UMAP visualization of gene expression of marker genes, including *Gata2, Lmo2, Klf1, Cited4, Hba-a1* and *Hbb-bh1* (top), and violinplot of gene expression of marker genes across different cell types on the Embryogenesis dataset (bottom).

While the cell-type transition matrix successfully captured the differentiation trajectory from “Blood progenitors” to “Erythroid” (Fig. 3a), several cell subtypes remain to be resolved in the Embryogenesis dataset^16^. Furthermore, we observed that “Blood progenitors 1” exhibited the highest descendancy probability toward “Blood progenitors 2”, which in turn showed the highest descendancy probability toward “Erythroid 1”, collectively outlining a developmental path (Blood progenitors 1→Blood progenitors 2→Erythroid 1→Erythroid 2→Erythroid 3) (Fig. 3d). The above developmental path is clearly consistent with the cell type transition trends inferred by CytoTRACEKernel^29^ of CellRank 2^13^ (Fig. 3e). This developmental path also reflects the temporal proportions of cell subtypes across different time points. Specifically, the cell subtype “Erythroid 3”, situated at the terminal end of the path, is predominantly enriched at the final time point (Fig. 3e). We displayed the expression distributions of several marker genes for mouse embryonic development across cell subtypes (Fig. 3f), which may facilitate further characterization and identification of these cell subtypes. For example, *Gata2* is a known marker gene of Hematopoietic progenitor cell^30-32^, *Lmo2* is a marker gene of Hematopoietic cell^16, 32^, *Klf1* is a marker gene of early erythroid cell^31, 32^, *Cited4* is a marker gene of erythrocyte^16, 32^, *Hba-a1* and *Hbb-bh1* is marker genes of erythroid cell^16, 32^.

To summarize, scMTG effectively infers transition matrices across time points and generates cell-type transition matrices, thereby enabling the elucidation of known cell-type differentiation processes and the discovery of novel cell subtypes.

### scMTG infers biologically meaningful transition relationships and fate dynamics

To assess the capability of scMTG in resolving temporal dynamics, we conducted a series of comparative analyses leveraging the low-dimensional cell representations and the inferred transition matrices. First, we created a UMAP plot of the Pancreas^17^ dataset based on the scMTG-derived cell representations, and annotated the plot with time point and cell type labels, enabling visual inspection of cell subpopulation separability (Fig. 4a-b). Notably, cells from distinct time points exhibit distinct distributions, and cells of different types are clearly separated in the UMAP plot.

**Fig. 4.**
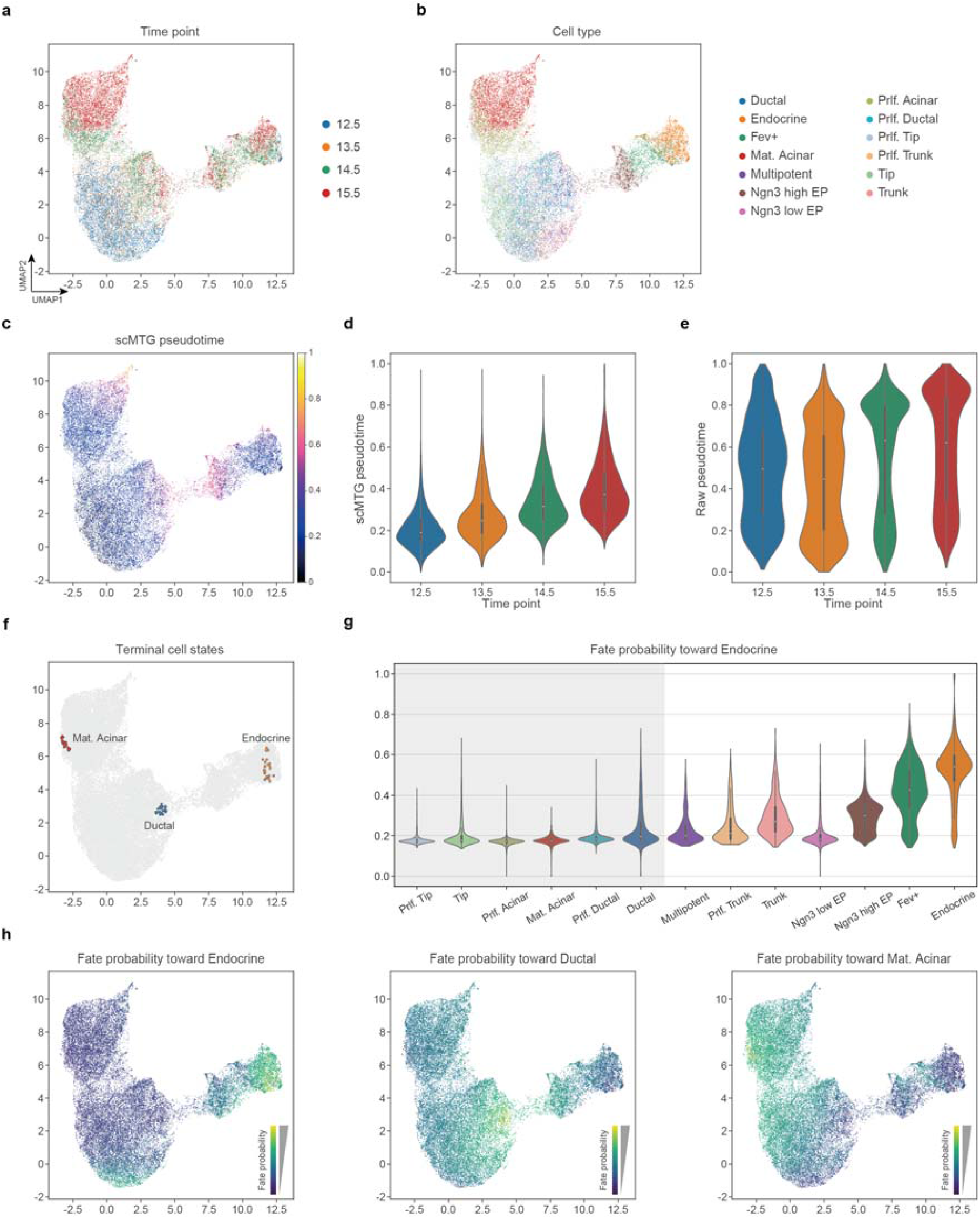
scMTG facilitates temporal analysis. **a-c**, UMAP visualization colored by time points (**a**), cell types (**b**) and inferred pseudotime based on scMTG embeddings (**c**) on the Pancreas dataset. **d**-**e**, Pseudotime distribution inferred based on scMTG embeddings (**d**) and raw data (**e**) across different time points on the Pancreas dataset. **f**, Terminal cell states identified from scMTG on the Pancreas dataset. **g**, Distribution of fate probabilities of different cell types toward the Endocrine terminal state on the Pancreas dataset. **h**, Cell fate probabilities toward the Endocrine (left), Ductal (middle) and Mat. Acinar (right) terminal states on the Pancreas dataset.

Second, we employed the CytoTRACEKernel^29^ of CellRank 2^13^ to infer pseudotime for the Pancreas^17^ dataset using different cell representations. The pseudotime inferred from the low-dimensional latent space representations of cells derived by scMTG was projected onto the aforementioned UMAP visualization (Fig. 4c), and showed a strong positive correlation with the experimentally annotated time points (Fig. 4d), yielding a Pearson correlation coefficient of 0.559. By comparison, the pseudotime inferred from the high-dimensional gene expression space representations of cells achieved a markedly lower correlation of 0.166 (Fig. 4e and Supplementary Fig. 5). These findings indicate that scMTG effectively preserves and enhances the temporal signal embedded in single-cell time-series data.

Third, we characterized the cell dynamics via random walks based on the transition matrices across time points inferred by scMTG, and identified three terminal states, including Endocrine cells, Ductal cells and Mature Acinar cells (Fig. 4f). These terminal states correspond to three well-characterized mature cell types in mouse pancreas development and closely align with the lineage relationship reported in the original study^17^ (Supplementary Fig. 6). To quantify developmental potential, we further computed fate probabilities toward each terminal state for all cells in the Pancreas^17^ dataset by quantifying the arrival frequencies from query cells to each terminal state over multiple random walks. Taking the Endocrine cells as an example, we examined how fate probabilities toward this terminal state varied across cell types, and observed a progressive increase along the developmental path to Endocrine cells, mirroring the gradual process of differentiation (Fig. 4g). Finally, we incorporated the cell fate probabilities toward all three terminal states onto the above UMAP plot (Fig. 4h), demonstrating that the cell fate probabilities are in strong agreement with the actual differentiation trajectories in mouse pancreas development (Supplementary Fig. 6).

Altogether, scMTG facilitates effective and interpretable temporal analysis, shedding light on the underlying temporal dynamics and cell fate decisions during development or differentiation.

### scMTG enables the construction of temporal GRNs

It is essential for understanding the mechanisms underlying cell fate decision and developmental differentiation to identify key genes that drive the transition processes and construct temporal GRNs from single-cell time-series data. Starting from the cell-type-specific transition gradient matrix, we first identified source genes whose downstream gradient profiles could be explained by the observed t1-to-t2 transcriptional shift. We then expanded the driver set using a residual-based procedure: after selecting the top source gene, additional genes were added only if their gradient profiles explained the remaining target-gene changes not captured by previously selected drivers. Finally, we constructed a cell-type-specific and transition-specific regulatory network by connecting selected source genes to target genes whose expression changes were directionally consistent with the corresponding gradient effects (see Methods and Fig. 5a-c).

**Fig. 5.**
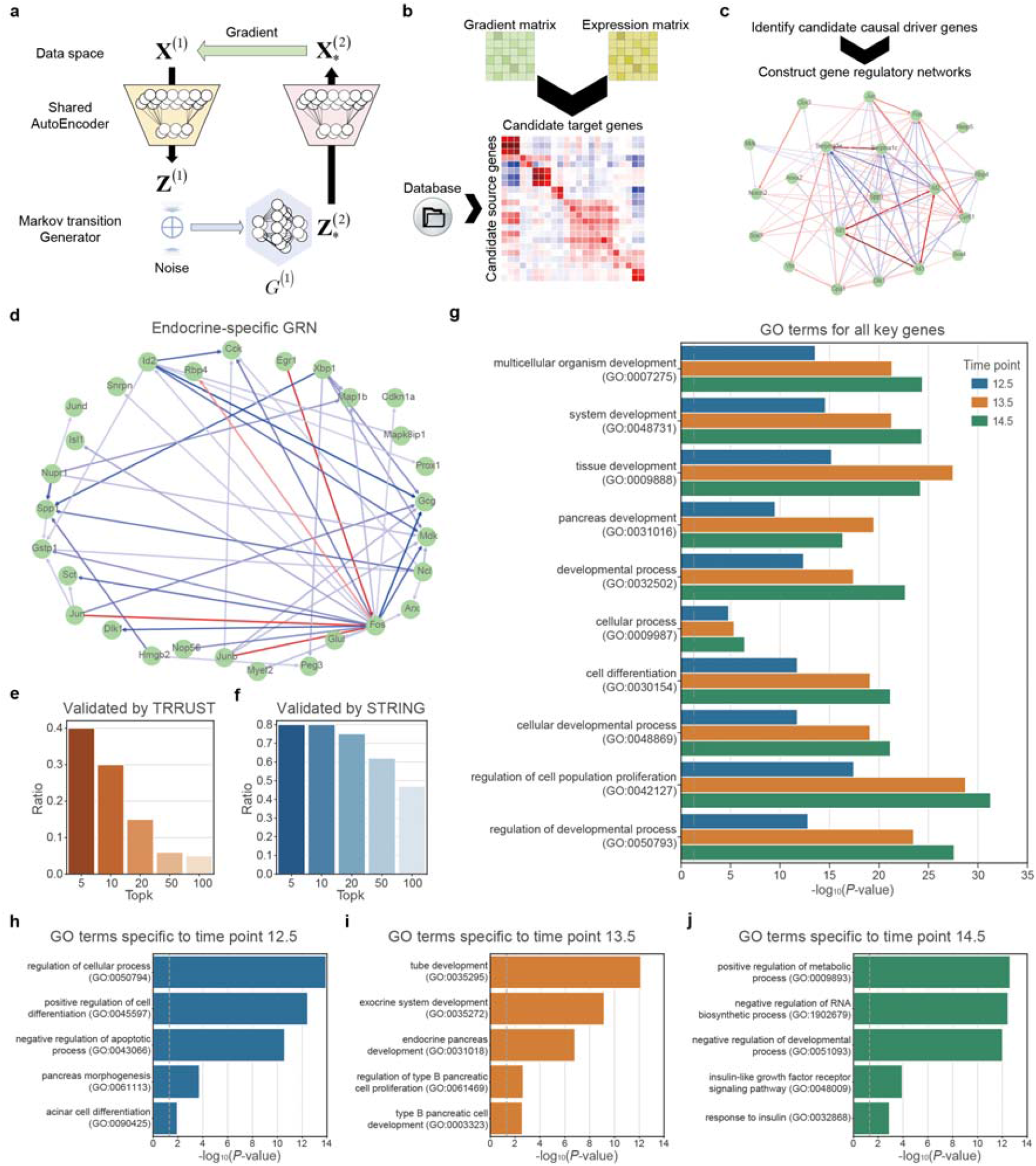
scMTG enables the construction of temporal GRNs. **a**, Generation of gradient matrix between adjacent time points. **b**, Selection of candidate source genes and candidate target genes from gradient matrix and expression matrix. **c**, Iterative identification of candidate causal driver genes and construction of transition-specific and cell-type-specific GRN. **d**, A showcase of Endocrine-specific GRN on the Pancreas dataset. **e-f**, In the Endocrine-specific GRN, the ratio of the top k regulatory relationships (key gene pairs) validated by TRRUST regulatory network database (**e**) and STRING protein-protein interaction network database (**f**). **g-j**, The GO terms obtained from all key genes (**g**) and specific key genes (**h-j**) of three time point pairs on the Pancreas dataset.

Specifically, we constructed multiple cell-type-specific temporal GRNs on the Pancreas^17^ dataset (Supplementary Fig. 7). For example, Endocrine-specific temporal GRN derived by scMTG contains cell-type-specific regulatory relationships and key genes, such as *Cck* and *Dlk1* reported in the previous literature^33, 34^ (Fig. 5d). To validate the biological significance of temporal GRNs, we compared the regulatory relationships within these GRNs against those derived from TRRUST regulatory network database^35^ and STRING protein-protein interaction network database^36^ (Fig. 5e-f). For example, among the top five regulatory relationships with the highest regulatory intensity in the Endocrine-specific GRNs, such as regulatory relationship between *Egr1* and *Fos* (Fig. 5e), two (40%) are validated by TRRUST database^35^, respectively. At least 47% of the top k regulatory relationships (k = 5, 10, 20, 50 and 100), as ranked by regulatory intensity, are validated by the STRING PPI database^36^. These outcomes demonstrate that scMTG can capture authentic gene regulatory relationships that have been supported by existing database.

Moreover, we constructed three temporal GRNs spanning adjacent time points on the Pancreas^17^ dataset comprising four time points (12.5, 13.5, 14.5 and 15.5) (Supplementary Fig. 8). These temporal GRNs consist of 123, 122, and 71 genes, respectively (Supplementary Fig. 9). Notably, 31 genes (25.2% to 43.7% overlapping rate) are shared across all three temporal GRNs, indicating that scMTG can learn general key genes across different pairs of time points. To further explore the biological processes represented by the shared or specific genes, we performed GO enrichment analysis on these distinct gene sets^37, 38^. The key genes of the above three temporal GRNs are jointly enriched in development-related or organ function-related GO terms such as *tissue development, developmental process* and *pancreas development*, cell regulation-related GO terms such as *regulation of cell population proliferation*, and cell development and differentiation-related GO terms such as *cellular developmental process* and *cell differentiation* (Fig. 5g). In contrast, the specific key genes in each of the three temporal GRNs are enriched in unique GO terms mirroring different biological processes (Fig. 5h-j). For example, the specific key genes in the first temporal GRN are enriched in GO terms related to organogenesis and cell proliferation, including *pancreas morphogenesis* and *acinar cell proliferation* (Fig. 5h), while those in the second temporal GRN are enriched in GO terms associated with secretory system, including *tube development, exocrine system development* and *endocrine pancreas development* (Fig. 5i), and those in the third temporal GRN are enriched in GO terms linked to metabolic process and insulin, including *positive regulation of metabolic process* and *response to insulin* (Fig. 5j). These findings suggest that scMTG is capable of revealing both common regulatory mechanisms underlying mouse pancreas development and stage-specific regulatory mechanisms at distinct developmental stages.

In summary, by constructing biologically meaningful and literature-supported temporal GRNs, scMTG provides a powerful framework for deciphering the molecular regulatory mechanisms that govern dynamic biological processes.

## Discussion

In this study, we present scMTG, a Markov transition-based generative framework designed for modeling cell state transitions and inferring dynamic gene regulation from single-cell time-series data. The central conceptual advance of scMTG is to formulate single-cell temporal analysis as learning explicit, differentiable transition functions between adjacent time points, rather than only estimating finite-sample couplings between observed cell populations. By leveraging parameter-shared autoencoder module and time-specific Markov transition generators, scMTG jointly learns a shared representation space and conditional generative transition functions that describe how cell states evolve over time. This formulation provides a unified framework for temporal interpolation, transition matrix inference, fate-related analysis and transition-function-derived regulatory network construction. Across multiple benchmark datasets, scMTG accurately recovered held-out intermediate time points, inferred biologically meaningful transition relationships, and revealed dynamic regulatory programs associated with cell fate decisions. These results support scMTG as a general framework for reconstructing cellular dynamics and connecting temporal state transitions with regulatory mechanism discovery.

We anticipate several future directions to improve our method. First, scMTG assumes a first-order Markov structure in which the future cell state depends primarily on the current state. This assumption provides a tractable and interpretable framework for modeling adjacent time-point transitions, but more complex biological processes may involve longer temporal memory, unobserved intermediate states or external environmental cues. Future extensions could incorporate higher-order temporal dependencies or continuous-time formulations to better model single cell time-series datasets. Second, the current implementation uses separate transition generators for adjacent time-point pairs. While this design provides flexibility for stage-specific dynamics, the number of generators increases with the number of time points. Developing shared or time-conditioned transition generators may improve scalability for long time-series datasets. Third, although the gradient-derived regulatory networks provide candidate transition-associated regulatory drivers, they should not be interpreted as strict causal regulatory networks. Experimental perturbation validation^39^ or rigorous causal modeling^40, 41^ remains necessary to establish causal regulatory relationships. Last, with the continuous advancement of single-cell time-series multi-omics sequencing and spatiotemporal omics sequencing technologies^42-44^, we expect scMTG to be extended to leverage data generated by these technologies for more comprehensive and integrative regulatory analysis of dynamic biological processes. Together, scMTG provides a general and interpretable framework for reconstructing cellular dynamics and uncovering the regulatory programs that shape development, differentiation and disease progression.

## Methods

### The scMTG framework

scMTG takes the processed cell-by-gene matrix 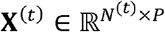 as input, where *N^(t)^* is the number of cells at time point *t*(*t* ∈ {1, ⋯, *T*}) and *P* is the number of genes, after a series of preprocessing operations including normalization, logarithmic transformation and selection of top 3000 variable genes using Scanpy^45^. scMTG is comprised of two key modules: an autoencoder module and a Markov transition module. The autoencoder module of scMTG shares parameters across all time points, enabling it to project cells from different time points into a shared latent space, thereby obtaining low-dimensional representations of cells 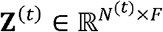, where *F* is the dimensionality of latent space.

We regard the biological process described by single-cell time-series data across multiple time points as a special type of stochastic process, and assume that the cell state at the current time point depends primarily on the previous time point. The central goal of the Markov transition module is therefore to learn a conditional transition kernel characterizes the distribution of future cell states given a source cell state at the current time point. In scMTG, we instantiate this transition kernel using a generative adversarial network (GAN)^46^. Specifically, for each pair of adjacent time points, the generator takes a source cell representation together with a random noise vector as input and outputs a simulated future cell state, thereby defining an explicit stochastic transition function. The discriminator encourages the generated future cell states to match the empirical cell-state distribution at the next time point. This design is particularly suitable for cell distribution match given unpaired single-cell time-series data. GANs have been previously adopted for distribution match of cell population in single cell setting^47, 48^. Importantly, the role of GANs here is not merely to generate realistic cells, but to learn an explicit continuous and differentiable transition function. This explicit feed-forward generator makes it straightforward to derive cell-to-cell transition probabilities and compute transition potential (e.g., gradient-based) for temporal regulatory network analysis. Although alternative conditional generative architectures, such as diffusion or flow-matching models, could in principle be incorporated into this framework^49-51^, the explicit differentiable generator used in scMTG provides a more direct implementation of the transition-kernel formulation.

For single-cell time-series data comprising *T* time points, this module consists of (*T*-1) GANs. Taking the *t*-th time point as an example, the generator *G*^(*t*)^ takes as input the concatenation of **Z**^(*t*)^ and a random noise term δ∼*N*(0,σ^2^), and outputs the simulated data at the (*t*+1)-th time point, denoted as 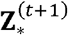:

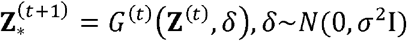

This induces the transition kernel ***K***_*t*_(*Z*^(*t+* 1)^|*Z*^(*t*)^) that maps the cell state distribution from time t to time t+1. The discriminator *D*(*t*) distinguishes whether ***Z***^(*t*+1)^ and 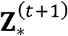 are real samples from the (*t*+1)-th time point or synthetic samples generated by *G*^(*t*)^.

### Model training

To more effectively optimize the different components of scMTG, we design a hybrid loss function consisting of several components.

First, similar to related studies^52, 53^, scMTG employs the Negative Binomial (NB) distribution to model single-cell time-series gene expression data:

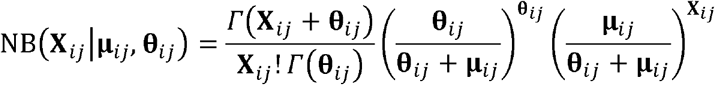

where **X**_*ij*_ is the gene expression value for the *i*-th cell and the *j*-th gene, **µ**_*ij*_ and **θ**_*ij*_ are mean and dispersion parameters of autoencoder module. Let *ℬ*_0_ be a mini-batch of data for training, we incorporate the reconstruction loss to optimize the autoencoder module:

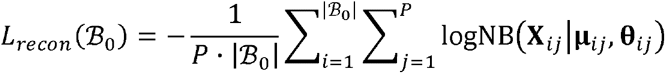

Second, we apply the generator loss and the loss of transition displacement to optimize the generators of Markov transition module:

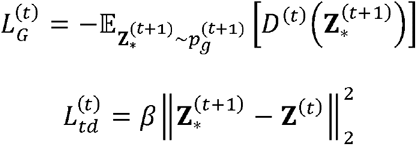

where 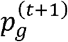 represents the distribution of generator between the *t*-th and (*t*+1)-th time points, and β is the weight of *L*_*td*_. Similar to Wasserstein distance used in optimal transport^7, 54^, *L*_*td*_ is designed under the assumption that cell state should not change too dramatically between adjacent time points.

Third, following the WGAN-GP^55^ framework, we employ the discriminator loss and the loss of gradient penalty to update the discriminators of Markov transition module:

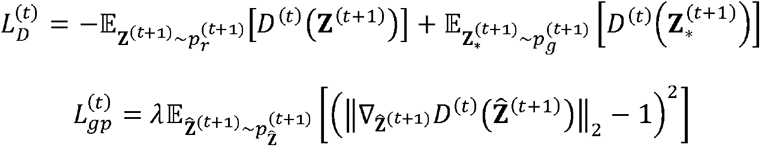

where 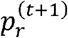 represents the distribution of real data at the (*t*+1)-th time point, 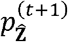 represents the distribution of sampled data 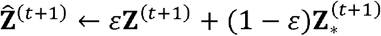 at the (*t*+1)-th time point, and λ is the weight of *L*_*gp*_. *L*_*gp*_ is introduced to stabilize the training of the GAN.

To enable joint training of the autoencoder module and Markov transition module, we iteratively update the parameters of the autoencoder (*AE*), generators (*G*), and discriminators (*D*) separately:

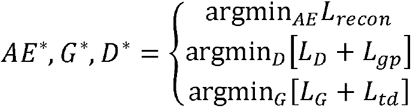

where *AE*\*, *G*\* and *D*\* denote the autoencoder, generators and discriminators obtained after the final training.

For all experiments, we used a four-layer fully-connected encoder network (2048-1024-512-256) and a four-layer decoder network with symmetric architecture, which is implemented with TensorFlow^56^. We employed the Adam stochastic optimization algorithm^57^ with a learning rate of 1e-4. For the datasets with <5 time points, mini-batch size for each time point is set to 32. For the datasets with >5 time points but <10 time points, mini-batch size is set to 16, and for the large number of time points with more than 10, mini-batch size is set to 8. Then we trained models with maximum iterations of 100,000 and early stopping when no reductions on training loss in 1000 iterations. For all experiments, we set the dimension of the latent embedding to 30 and trained models with β of 5, and λ of 10.

### Cell-specific growth-weighted sampling

To account for cell proliferation and apoptosis^7, 18, 19^, we adopt an estimated cell-specific growth-weighted sampling strategy during the training process. We first calculate the mean of the z-scores of genes annotated to proliferation and apoptosis^58-62^, as the birth score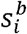 and death score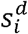 for the *i*-th cell. Then, the birth and death rates are estimated using the corresponding birth and death scores:

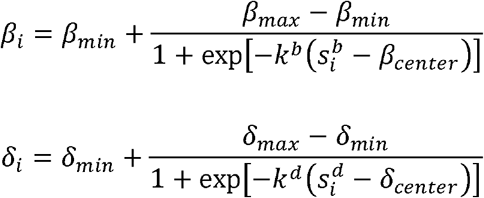

Where *β*_*min*_, *β*_*max*_, *β*_*center*_ and *k*^*b*^ *δ*_*min*_, *δ*_*max*_, *δ*_*center*_ and *k*^*d*^) denote the default parameters for birth process (death process). We ultimately obtain the growth rate for the *i*-th cell, which is used as the probability of sampling this cell during training after normalization.

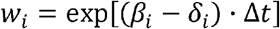

where Δ*t* denote the specific time interval between adjacent time points.

### Model evaluation

To validate the advantages of scMTG in interpolation, we compared scMTG to multiple baseline methods in this study, including WOT (https://github.com/broadinstitute/wot)^7^, PRESCIENT (https://github.com/gifford-lab/prescient)^19^, CellRank 2 (https://github.com/theislab/cellrank)^13^ and moscot (https://moscot-tools.org).^18^

Specifically, we first excluded cells from the intermediate time point designated for interpolation from the training data, and then interpolated single-cell data of held-out time points in the latent space using the transition matrix bridging the two adjacent time points. With the exception of PRESCIENT, which performs interpolation via a learned potential function, all other methods computed the transition matrix between the two adjacent time points flanking the held-out time points and subsequently performed interpolation within the latent space. To ensure a fair comparison, the latent-space interpolations generated by each respective method were projected back into the original feature space for the assessment of interpolation. Finally, we employed maximum mean discrepancy (MMD)^23, 24^, Spearman’s correlation coefficient (SCC)^18^, and local inverse Simpson’s index (LISI)^25^ as evaluation metrics. MMD quantifies the distance between the distributions of real and generated data, with smaller values indicating better generation performance. SCC measures the correlation between real and generated data, with larger values signifying superior generation quality. LISI evaluates the batch effect between the distributions of real and generated data, with larger values denoting better batch mixing.

### Generation of transition matrix

We represent the transition matrix from the *t*-th time point to the (*t*+1)-th time point as 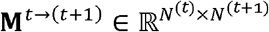,where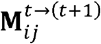 denotes the transition probability from the *i*-th cell at the *t*-th time point to the *j*-th cell at the (*t*+1)-th time point. To generate the transition matrix M^*t* →(*t*+1)^ we employed a three-step procedure.

First, the generator from the *t*-th time point to the (*t*+1)-th time point within the Markov transition module of scMTG receives concatenated input comprising the low-dimensional representations of cells the *t*-th time point and random noise, and derives the generated cells at the (*t*+1)-th time point. To produce a stable transition matrix, we introduced *N*_*noise*_ (default 1000) distinct random noise vectors for each cell representation, thereby obtaining *N*_*noise*_ generated cells at the (*t*+1)-th time point through the generator.

Second, for the *i*-th cell at the *t*-th time point, based on the *N*_*noise*_ cells generated for the (*t*+1)-th time point, we constructed a Gaussian mixture model (GMM) composed of *N*_*noise*_ Gaussian distributions with mean of 0 and standard deviation of 0.5. For the *j*-th cell at the (*t*+1)-th time point, we considered its probability density under this GMM as the transition probability. As a result, we derived the transition probabilities from each cell at the *t*-th time point to each cell at the (*t*+1)-th time point, yielding the initial transition matrix.

Third, we row-normalized the initial transition matrix to ensure valid transition probabilities and applied an adaptive thresholding strategy to sparsify the matrix^13^, thereby acquiring the final transition matrix **M**^*t*→*t*+1)^.

### Accuracy metrics at the germ-layer and cell-type levels

To quantitatively evaluate the performance of transition matrix, we introduced two evaluation metrics defined at the germ-layer and cell-type levels. First, we mapped each cell type to its corresponding germ layer, such as neuroectoderm, surface ectoderm, endoderm, mesoderm and extraembryonic, and excluded those that could not be unambiguously assigned to a specific germ layer. Motivated by the biological constraint that cells rarely cross germ-layer boundaries^28^, we filtered out the transitions with edge weights below 0.05, and considered transitions occurring within the same germ layer as correct, whereas those crossing germ-layer boundaries were treated as incorrect. The germ-layer transition accuracy was calculated as the ratio of the weighted sum of transitions satisfying the respective germ-layer boundaries to the weighted sum of all evaluated transitions, with edge weights serving as the weighting factors.

Second, we benchmarked each predicted transition against a curated reference set of biologically allowed transitions (Supplementary Table 2). We constructed this reference set by systematically surveying the literature for all cell types in the Embryogenesis dataset^16^ and documenting their established ancestral and descendant cell types during mouse embryogenesis. The cell-type transition accuracy was calculated as the ratio of the weighted sum of transitions satisfying the respective cell-type restrictions to the weighted sum of all evaluated transitions, with edge weights serving as the weighting factors.

### GRN construction of scMTG

The process for constructing temporal GRNs is illustrated in Fig. 5a-c. Let *c* denote the cell type of interest. Let 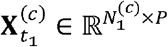 denote the expression matrix for cells of type *c* at time *t*_*1*_, and let 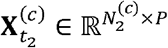 denote the expression matrix for cells of type *c* at time *t*_*2*_. Here, *P* is the number of genes after preprocessing.

For each source cell *m* at time *t*_*1*_, let **G**^(*m*)^ ∈ ℝ^*P*x*P*^ denote the gradient matrix learned from scMTG. The entry 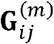 measures the total transition effect from source gene *i* at *t*_*1*_ to target gene *j* at *t*_*2*_:

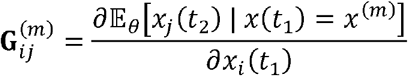

For the selected cell type *c*, we define the cell-type-specific average gradient matrix as:

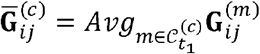

Where 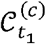 denotes the set of cells of type *c* observed at time *t*_*1*_. In practice, we use the median function as *Avg*.

### Step 1. Define the cell-type-specific transition direction

We first computed the average expression change from *t*_*1*_ to *t*_*2*_ within cell type *c*:

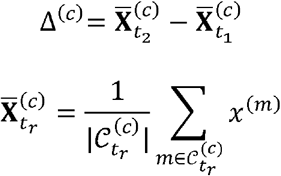

Here,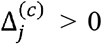 indicates that target gene *j* is up-regulated from *t*_*1*_ to *t*_*2*_ in the cell type of interest, whereas 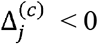 indicates down-regulation.

### Step 2. Select candidate target genes

Target genes are genes whose *t*_*1*_-to-*t*_*2*_ changes should be explained by candidate source drivers. Let *P* denote the target-gene set. We computed the incoming gradient strength for each candidate target gene *j*:

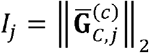

where *C* is the set of candidate source genes defined below, and 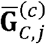 denotes the column of 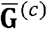 restricted to source genes in *C*. We then defined *P* as:

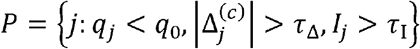

where *q*_*0*_ is the adjusted p-value threshold by comparing profile at time *t*_*1*_ and *t*_*2*_. In practice, we set *q*_*0*_=0.05, *τ* _Δ_=0.02, and *τ*_I_=0.02 in default.

We assigned each target gene a transition weight:]

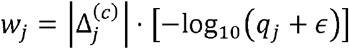

The weight *w*_*j*_ is large when gene *j* is strongly and significantly changed during the transition.

### Step 3. Select candidate source genes

We restricted the candidate set *C* to biologically plausible regulators^35, 63, 64^:

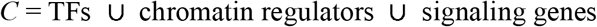

### Step 4. Compute a transition-driving score for each source gene

For each source gene *i*∈*C*, we defined its downstream gradient profile over the target genes *P* as:

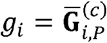

We then defined the weighted target transition vector and the weighted gene-specific gradient profile:

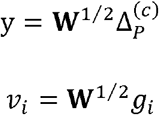

where

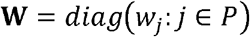

A simple direction-aware transition-driving score is:

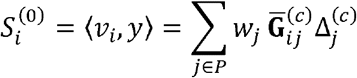

A source gene *i* is prioritized if its downstream gradient profile is aligned with the observed *t*_*1*_-to-*t*_*2*_ transcriptional shift. This score is more informative than ranking genes by raw gradient magnitude alone.

### Step 5. Identifying a candidate causal driver gene set

We first selected the strongest initial source driver:

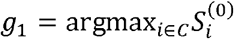

The selected driver set is initialized as:

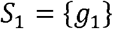

At iteration *k*, we fitted non-negative coefficients for the currently selected drivers:

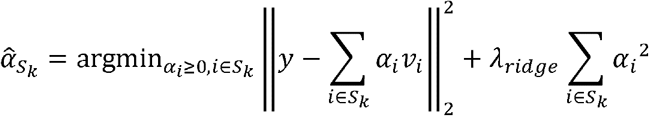

where the regularization weight *λ*_*ridge*_ defaults to 0.1. Essentially, we expected the gradient profiles of selected driver genes can decompose and explain the observed transcriptional changes of target genes.

We then computed the unexplained transition residual:

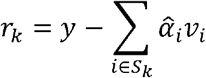

The residual *r*_*k*_ represents the component of the target-gene transition that is not explained by the currently selected drivers.

Next, we selected the source gene whose gradient profile best explains the remaining residual:

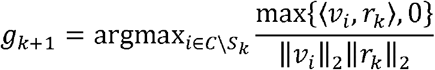

The selected driver set is then updated as:

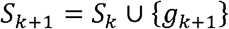

This step adds a new source gene only if its gradient profile explains a component of the target-gene transition that is not already captured by the existing selected drivers. This residual-based expansion reduces redundancy among co-expressed marker genes and favors complementary transition-driving candidates.

Stop when residual improvement is small (less than 0.0001), the driver set is sufficiently small.

We kept driver *i* only if its gradient profile is aligned with the observed transition (Directional Consistency):

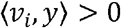

### Step 6. Construct a transition-specific and cell-type-specific GRN

After selecting the driver set *S*, we constructed a directed network from selected source drivers to target genes. For *i*∈*S* and *j*∈*P*, the direction-consistent edge weight is defined as:

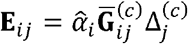

An edge *i*→*j* is retained if it satisfies all or most of the following criteria.

- The gradient magnitude is sufficiently large:

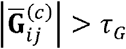

where *τ*_*G*_ defaults to 0.0001.
- The gradient direction is consistent with the target-gene change:

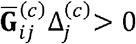

The resulting edge *i*→*j* means that, for the selected cell type and transition, source gene *i* is a selected transition-driver candidate and its gradient-supported effect on target gene *j* is directionally consistent with the observed *t*_*1*_-to-*t*_*2*_ change. Note that this is not a claim of direct causal regulation without further assumption.

## Data availability

All data used in this paper are publicly available via the original publications and releases. Pancreas dataset was downloaded in Gene Expression Omnibus (GEO) under accession number GSE132188^17^. CerebralCortex dataset was downloaded in GEO under accession number GSE162170^14^. Veres1 and Veres2 datasets were downloaded in GEO under accession number GSE114412^15^. Serum dataset was downloaded at https://doi.org/10.6084/m9.figshare.200514927, ^13^. Embryogenesis dataset was downloaded at https://ftp.ebi.ac.uk/biostudies/fire/E-MTAB-/967/E-MTAB-6967/Files/atlas_data.tar.gz^16^.

## Code availability

The scMTG software, including detailed documents and tutorial, is freely available on GitHub (https://github.com/liuq-lab/scMTG).

## Acknowledgements

This work was supported by the National Key Research and Development Program of China grant nos. 2021YFF1200902 (R.J.), 2023YFF1204802 (R.J.), the National Natural Science Foundation of China grants nos. 62273194 (R.J.).

## Author contributions

Q.L., W.H.W and R.J. conceived the study and supervised the project. X.C. and Q.L. designed, implemented and validated scMTG. X.C. and Q.L. wrote the manuscript.

