## Supplementary File for "scMTG reconstructs single-cell temporal dynamics with Markov transition generators"

**Supplementary Figures**

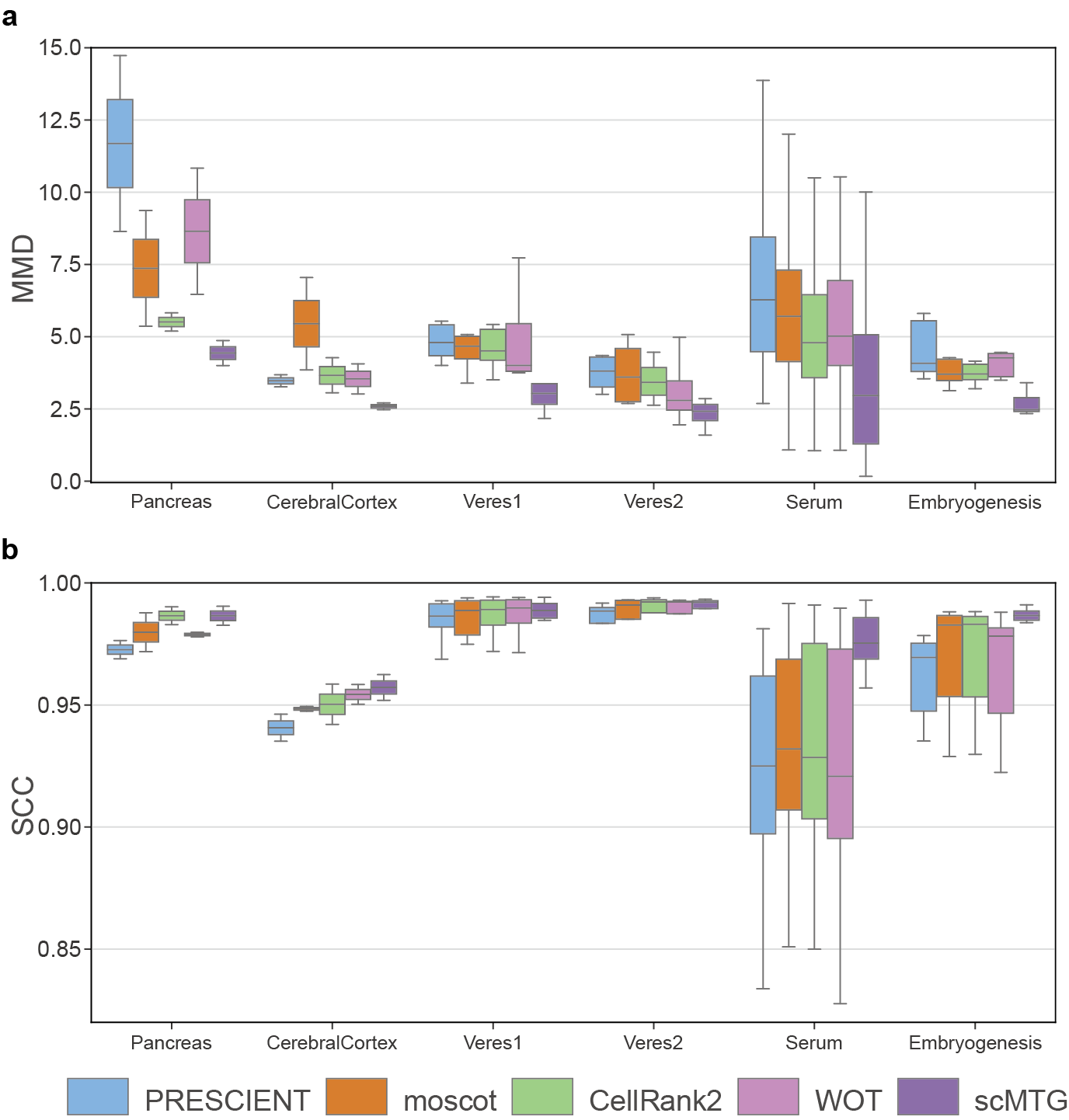

**Supplementary Fig. 1. Performance of interpolation**. **a-b**, Interpolation results of different methods evaluated by MMD (**a**) and SCC (**b**) on 6 benchmark datasets.

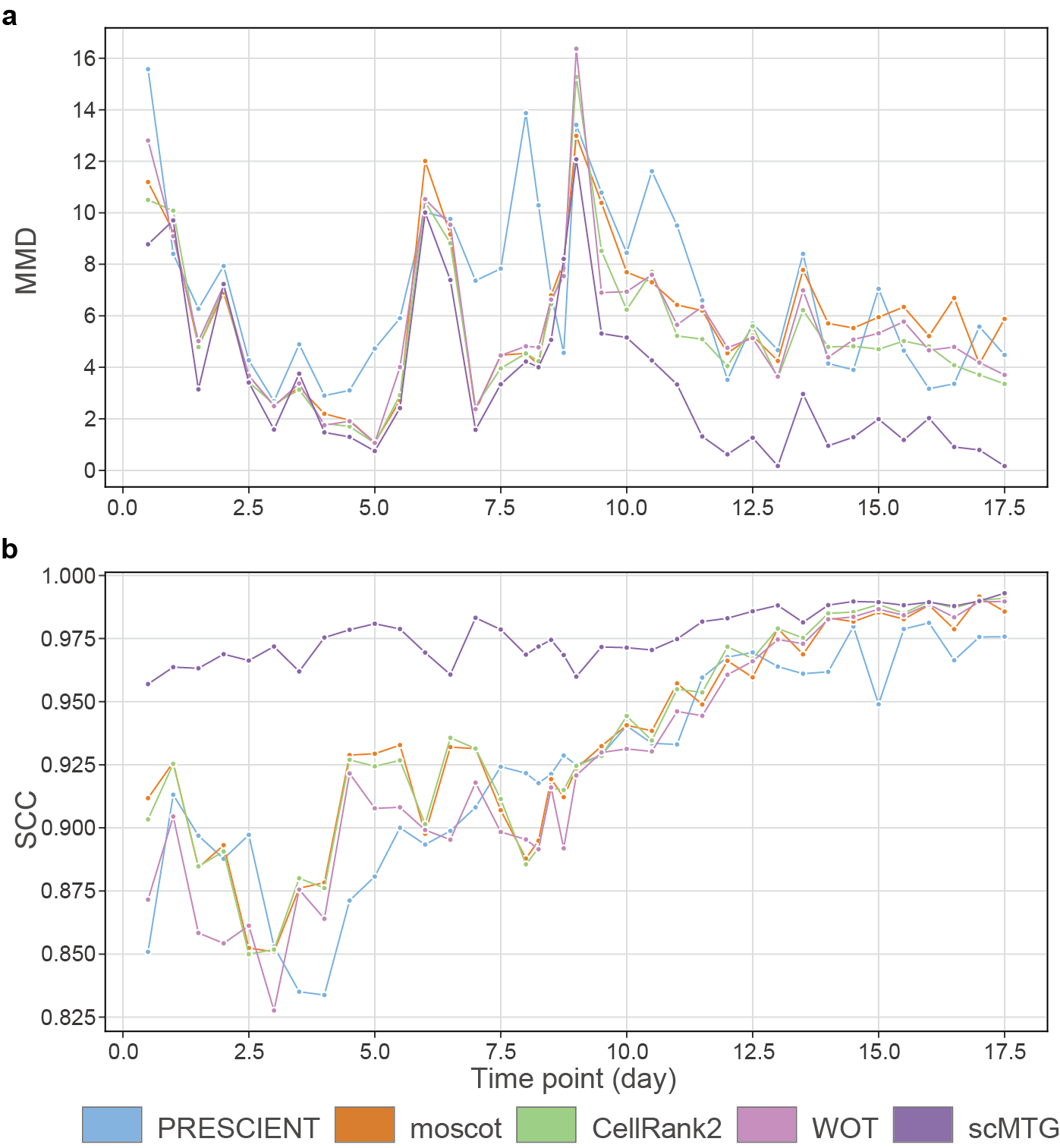

**Supplementary Fig. 2. Performance of interpolation on the Serum dataset**. **a-b**, Interpolation results of different methods across time points evaluated by MMD (**a**) and SCC (**b**) on the Serum dataset.

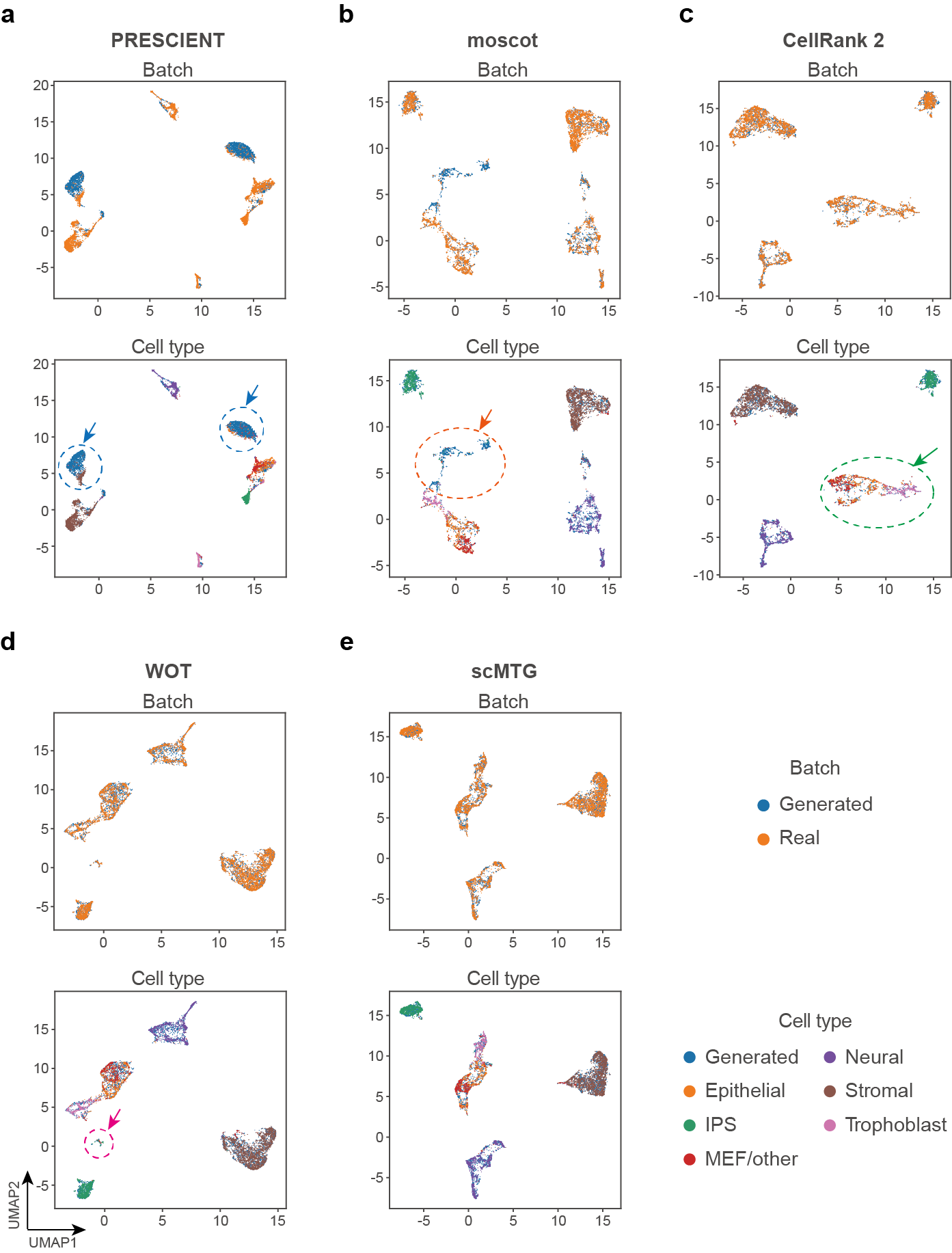

**Supplementary Fig. 3. UMAP visualization of the generated and real cells** **on the Serum dataset**. **a-e**, UMAP visualization of the real cells and generated cells derived from different methods, including PRESCIENT (**a**), moscot (**b**), CellRank 2 (**c**), WOT (**d**), and scMTG (**e**). For each method, cells are colored by data batch (top) and cell type (bottom).

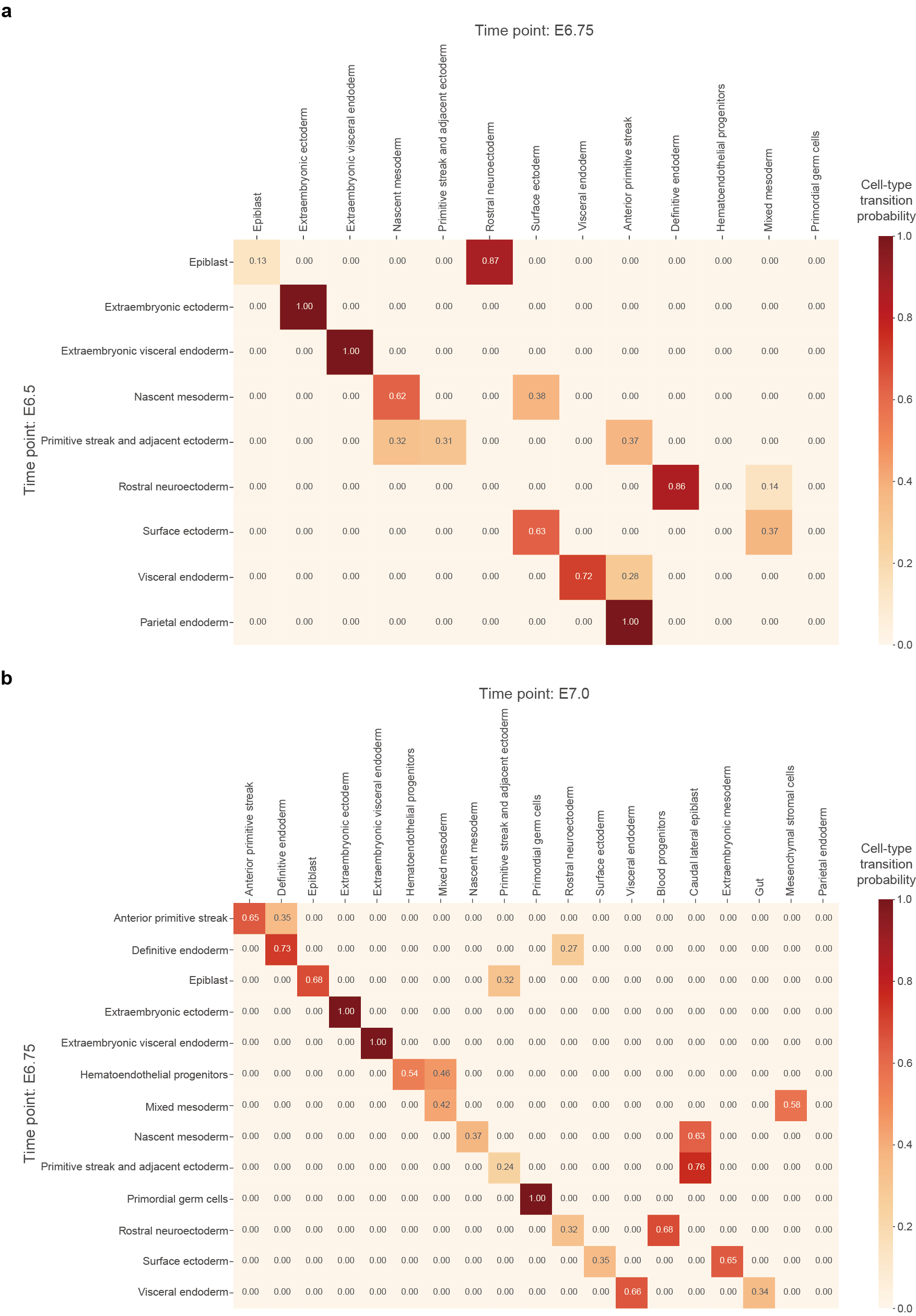

**Supplementary Fig. 4. Heatmap visualizing transition probabilities of cell types.** **a-g**, Heatmap visualizing transition probabilities of cell types from E6.5 to E6.75 (**a**), from E6.75 to E7.0 (**b**), from E7.0 to E7.25 (**c**), from E7.25 to E7.5 (**d**), from E7.5 to E7.75 (**e**), from E7.75 to E8.0 (**f**) and from E8.0 to E8.25 (**g**) on the Embryogenesis dataset as obtained using scMTG.

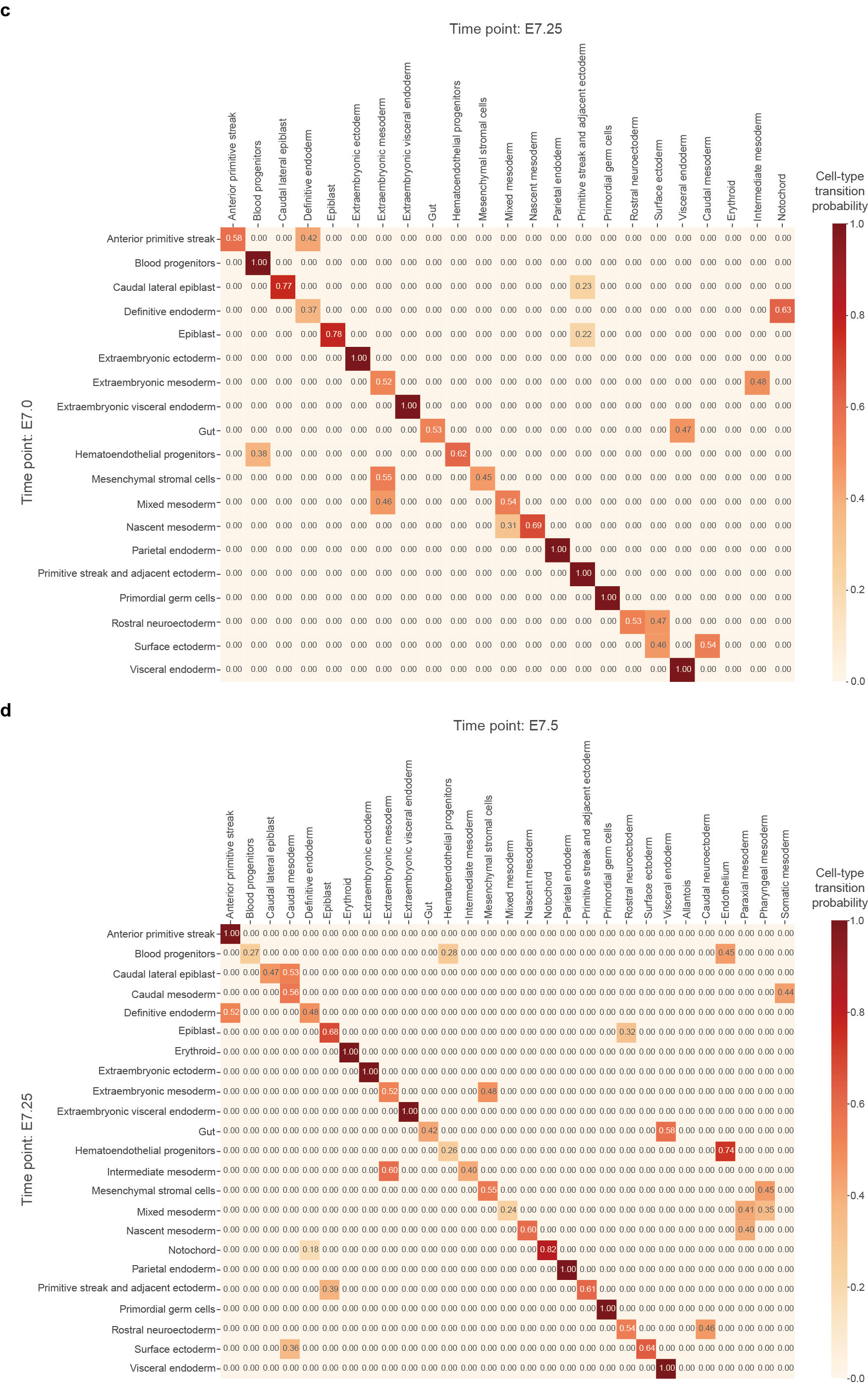

**Supplementary Fig. 4 (continue). Heatmap visualizing transition probabilities of cell types.** **a-g**, Heatmap visualizing transition probabilities of cell types from E6.5 to E6.75 (**a**), from E6.75 to E7.0 (**b**), from E7.0 to E7.25 (**c**), from E7.25 to E7.5 (**d**), from E7.5 to E7.75 (**e**), from E7.75 to E8.0 (**f**) and from E8.0 to E8.25 (**g**) on the Embryogenesis dataset as obtained using scMTG.

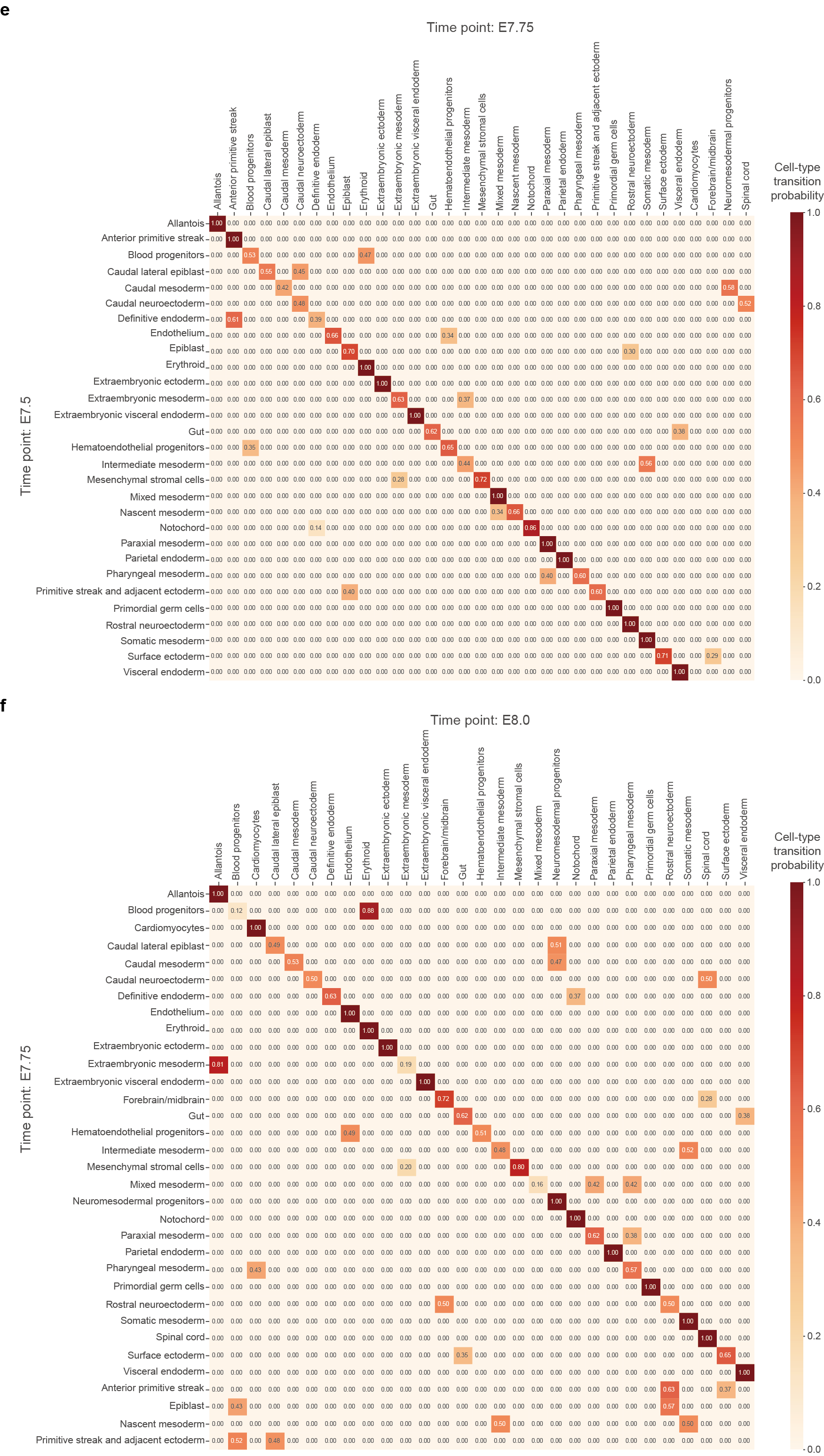

**Supplementary Fig. 4 (continue). Heatmap visualizing transition probabilities of cell types.** **a-g**, Heatmap visualizing transition probabilities of cell types from E6.5 to E6.75 (**a**), from E6.75 to E7.0 (**b**), from E7.0 to E7.25 (**c**), from E7.25 to E7.5 (**d**), from E7.5 to E7.75 (**e**), from E7.75 to E8.0 (**f**) and from E8.0 to E8.25 (**g**) on the Embryogenesis dataset as obtained using scMTG.

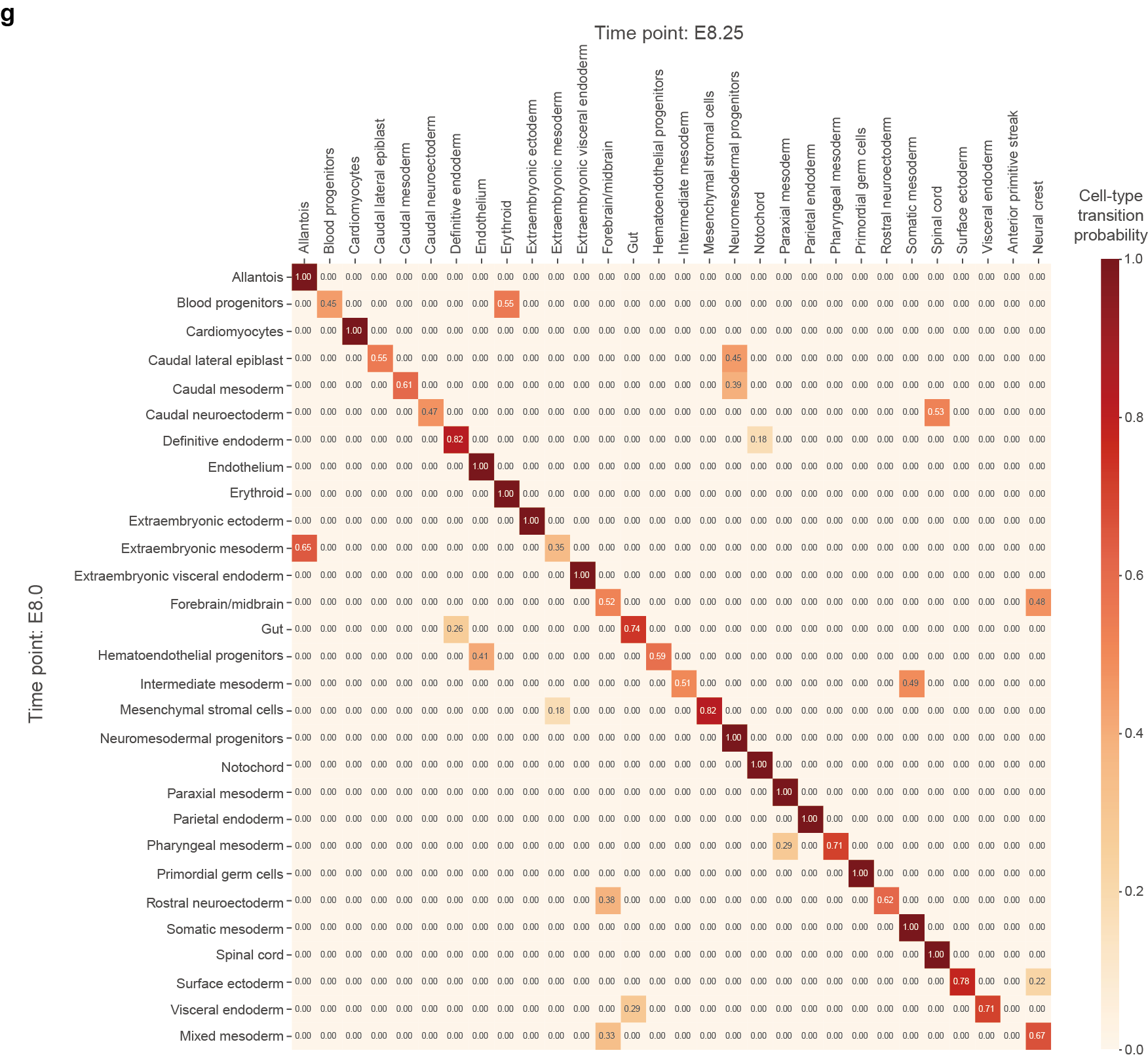

**Supplementary Fig. 4 (continue). Heatmap visualizing transition probabilities of cell types.** **a-g**, Heatmap visualizing transition probabilities of cell types from E6.5 to E6.75 (**a**), from E6.75 to E7.0 (**b**), from E7.0 to E7.25 (**c**), from E7.25 to E7.5 (**d**), from E7.5 to E7.75 (**e**), from E7.75 to E8.0 (**f**) and from E8.0 to E8.25 (**g**) on the Embryogenesis dataset as obtained using scMTG.

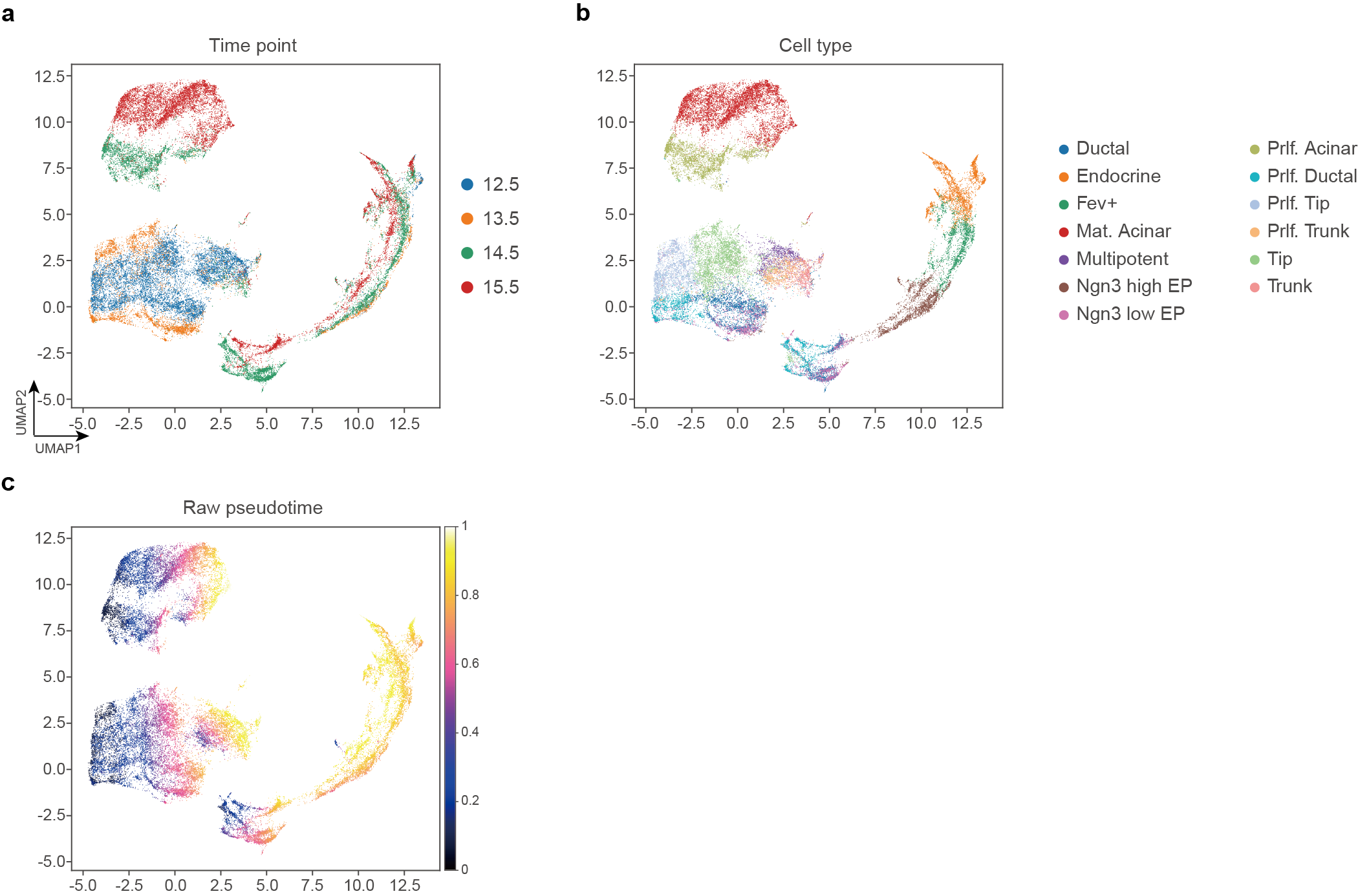

**Supplementary Fig. 5. UMAP visualization of raw data.** **a-c**, UMAP visualization colored by time points (**a**), cell types (**b**) and inferred pseudotime (**c**) based on raw data of the Pancreas dataset.

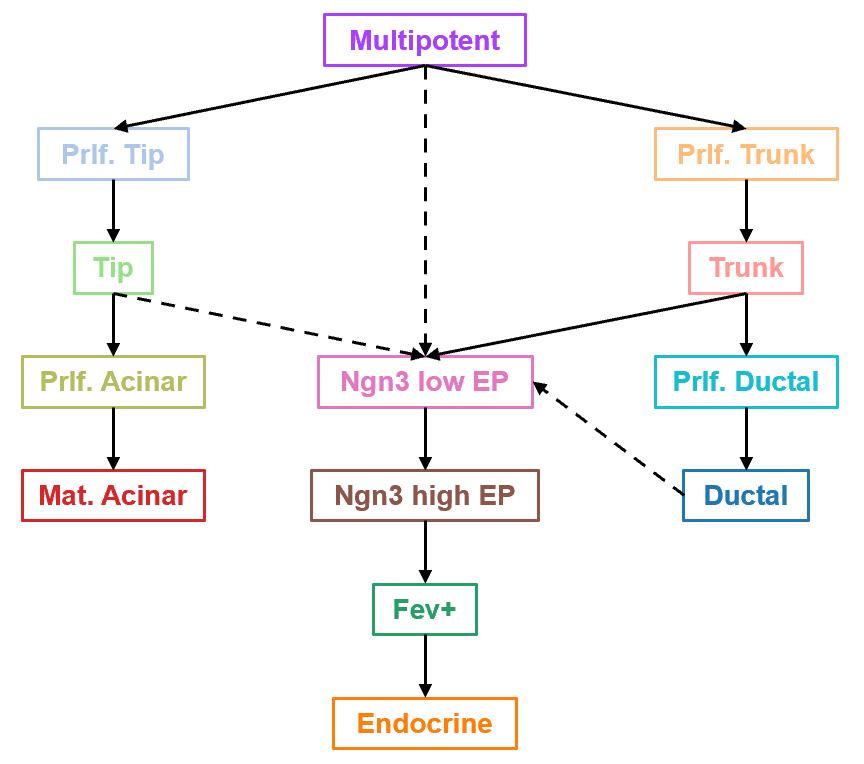

**Supplementary Fig. 6. Diagram of the lineage relationship between pancreatic cells at early embryonic stages.**

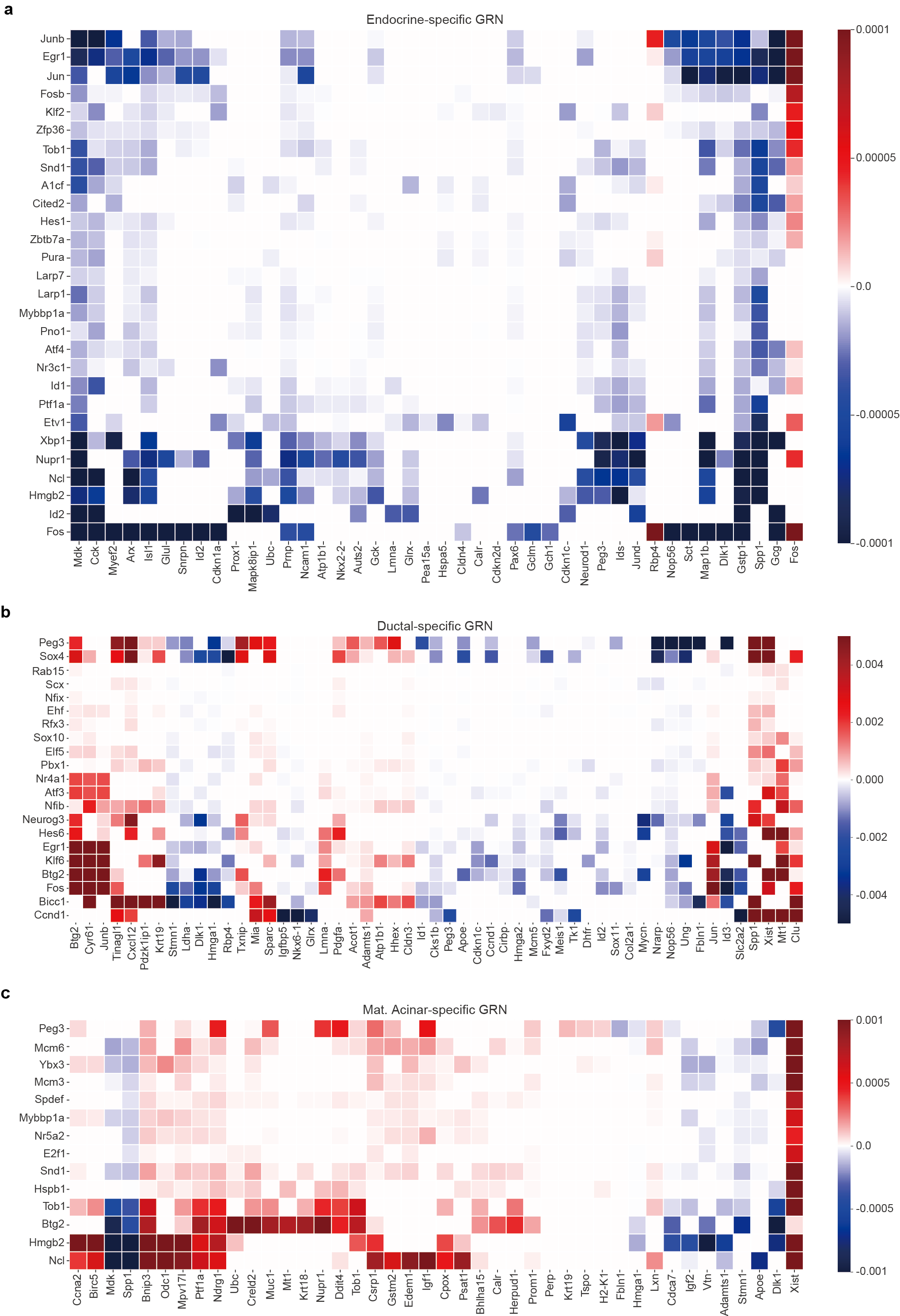

**Supplementary Fig. 7. Cell-type-specific temporal GRNs of the Pancreas dataset.** **a-c**, Heatmap of the matrix for regulatory relationships for Endocrine (**a**), Ductal (**b**) and Mat. Acinar (**c**) on the Pancreas dataset.

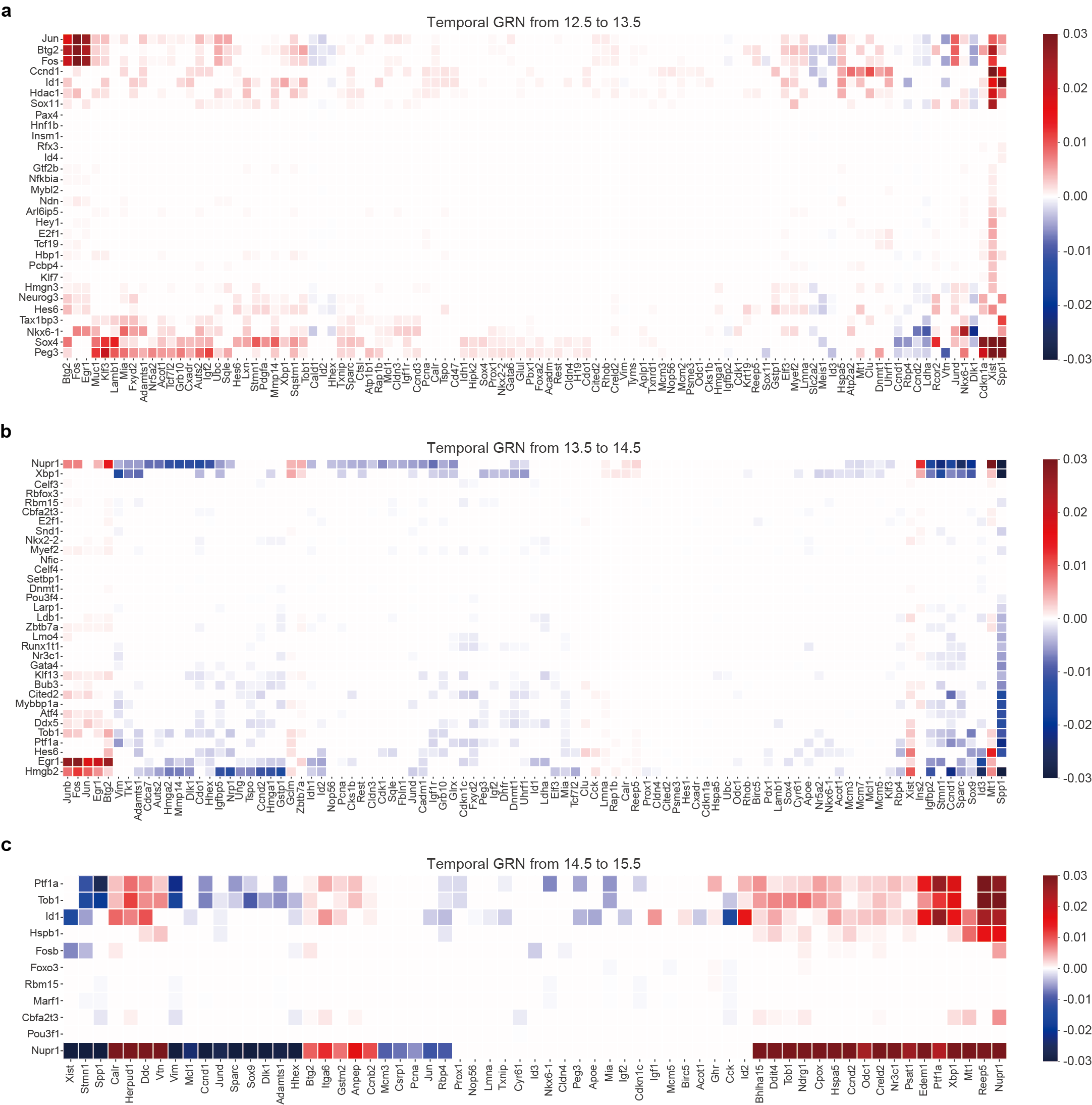

**Supplementary Fig. 8. Temporal GRNs of the Pancreas dataset.** **a-c**, Heatmap of the matrix for regulatory relationships from 12.5 to 13.5 (**a**), from 13.5 to 14.5 (**b**) and from 14.5 to 15.5 (**c**) on the Pancreas dataset.

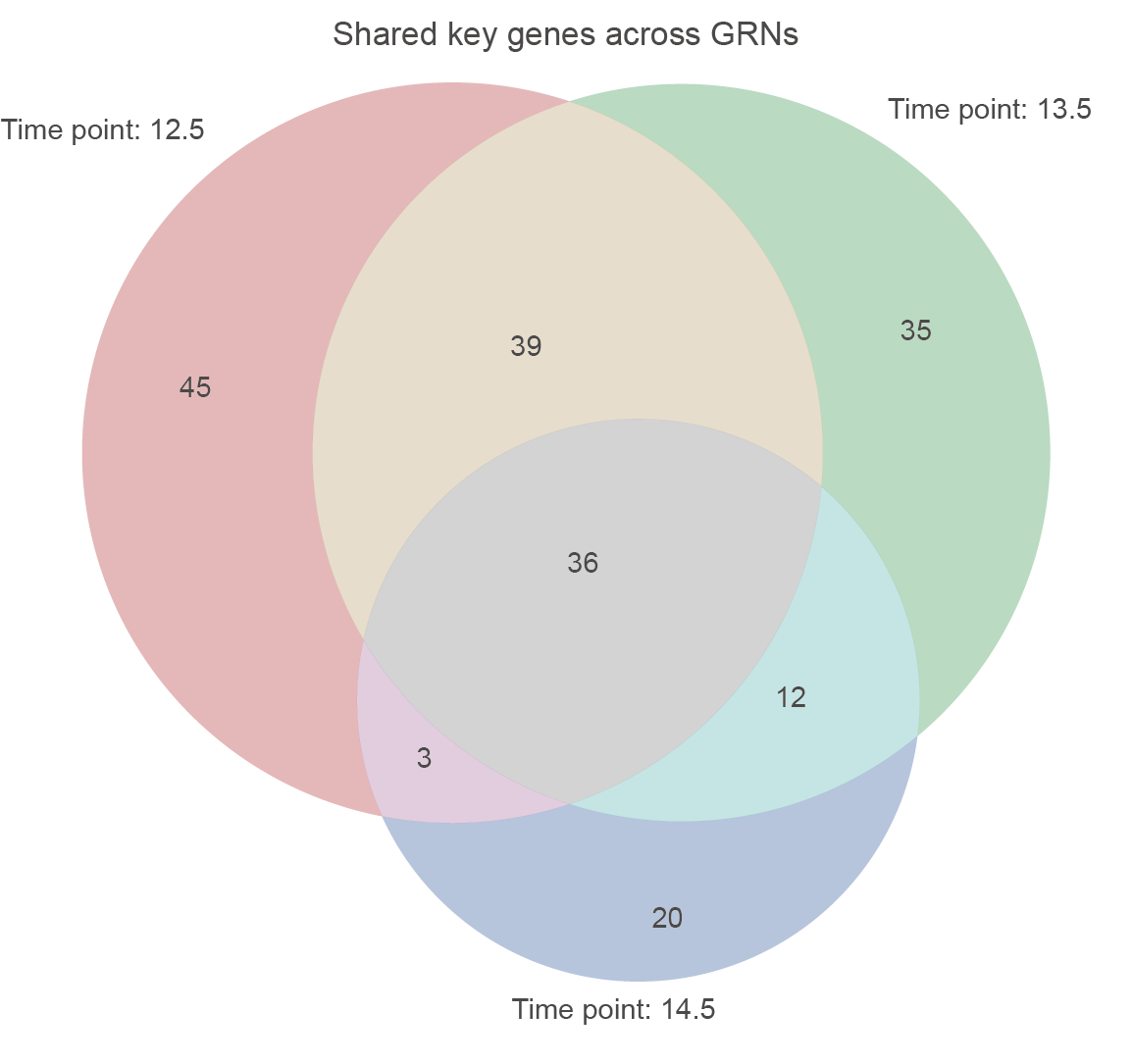

**Supplementary Fig. 9. Temporal GRNs with shared genes on the Pancreas dataset.** Venn diagram of key genes across three time point pairs identified by scMTG on the Pancreas dataset.

**Supplementary Tables**

**Supplementary Table 1. Summary of datasets used in this study.**

| **Dataset** | **Species** | **No. of cells** | **No. of genes** | **No. of cell types** | **No. of time points** |
| --- | --- | --- | --- | --- | --- |
| Pancreas^1^ | Mouse | 36,351 | 17,327 | 13 | 4 |
| CerebralCortex^2^ | Human | 57,868 | 33,355 | 23 | 4 |
| Veres1^3^ | Human | 51,274 | 16,224 | 12 | 8 |
| Veres2^3^ | Human | 38,494 | 16,039 | 8 | 6 |
| Serum^4, 5^ | Mouse | 165,892 | 19,089 | 7 | 39 |
| Embryogenesis^6^ | Mouse | 117,820 | 29,452 | 37 | 9 |
